# *Bacillus* endospore appendages form a novel family of disulfide-linked pili

**DOI:** 10.1101/2020.08.21.260141

**Authors:** Brajabandhu Pradhan, Janine Liedtke, Mike Sleutel, Toril Lindbäck, Ann-Katrin Llarena, Ola Brynildsrud, Marina Aspholm, Han Remaut

## Abstract

*Bacillus cereus sensu lato* is a group of Gram-positive endospore-forming bacteria with high ecological diversity. Their endospores are decorated with micrometer-long appendages of unknown identity and function. Here we isolate endospore appendages (Enas) from the food poisoning outbreak strain *B. cereus* NVH 0075-95 and find proteinaceous fibers of two main morphologies. By using cryo-EM and 3D helical reconstruction we show that *Bacillus* Enas form a novel class of Gram-positive pili. Enas consist of single domain subunits with jellyroll topology that are laterally stacked by β-sheet augmentation. Enas are longitudinally stabilized by disulfide bonding through N-terminal connector peptides that bridge the helical turns. Together, this results in flexible pili that are highly resistant to heat, drought and chemical damage. Phylogenomic analysis reveals the presence of defined *ena* clades amongst different eco- and pathotypes. We propose Enas to represent a novel class of pili specifically adapted to the harsh conditions encountered by bacterial spores.

## Introduction

When faced with adverse growth conditions, bacteria belonging to the phylum *Firmicutes* can differentiate into the metabolically dormant endospore state. These endospores exhibit extreme resilience towards environmental stressors due to their dehydrated state and unique multilayered cellular structure, and can germinate into the metabolically active and replicating vegetative growth state even hundreds of years after their formation (Setlow, 2014). In this way, Firmicutes belonging to the classes Bacilli and Clostridia can withstand long periods of drought, starvation, high oxygen or antibiotic stress.

Endospores typically consist of an innermost dehydrated core which contains the bacterial DNA. The core is enclosed by an inner membrane surrounded by a thin layer of peptidoglycan that will become the cell wall of the vegetative cell that emerges during endospore germination. Then follows a thick cortex layer of modified peptidoglycan that is essential for dormancy. The cortex layer is in turn surrounded by several proteinaceous coat layers (Atrih and Foster, 1999). In some *Clostridium* and most *Bacillus cereus* group species, the spore is enclosed by an outermost loose-fitting paracrystalline exosporium layer consisting of (glyco)proteins and lipids (Stewart, 2015). The surface of *Bacillus* and *Clostridium* endospores can also be decorated with multiple micrometers long and a few nanometers wide filamentous appendages, which show a great structural diversity between strains and species (Hachisuka and Kuno, 1976; Rode et al., 1971; Walker et al., 2007). Spores of species belonging to the *B. cereus* group are often covered with appendages which morphologically resemble pili of Gram-negative and Gram-positive bacteria when imaged by negative strain TEM (Ankolekar and Labbe, 2010; Smirnova et al., 2013). The endospore appendages, hereafter called Enas, vary in number and morphology between *B. cereus* group strains and species, and some strains even simultaneously express Enas of different morphologies (Smirnova et al., 2013). Structures resembling the Enas have not been observed on the surface of the vegetative cells suggesting that they may represent spore-specific fibers.

Although the presence of endospore appendages in species belonging to the *B. cereus* group was reported already in the ‘60s, efforts to characterize their composition and genetic identity have failed due to difficulties to solubilize and enzymatically digest the fibers (DesRosier and Lara, 1981; Gerhardt and Ribi, 1964). Therefore, there is no genetic or structural information and very limited functional data available for endospore appendages.

Here we isolate Enas from the food poisoning outbreak strain *B. cereus* NVH 0075-95 and find proteinaceous fibers of two main morphologies. By using cryo-EM and 3D helical reconstruction we show that the major form of Enas represents a novel class of Gram-positive pili. A unique architecture of subunit stabilization by lateral β-augmentation and longitudinal disulfide crosslinking gives rise to pili that combine high flexibility with high resistance to heat, drought and chemical damage. The genetic identity of the S-type Enas was deduced from the structural model and confirmed by analysis of mutants lacking genes encoding potential Ena protein subunits. S-type Ena fibers are encoded by three associated genes which are present in most species of the *B. cereus* group. Remarkably, recombinant Ena subunits spontaneously self-assemble *in vitro* and *in vivo* into protein nanofibers with native Ena-like properties and structure.

## Results

### *Bacillus cereus* NVH 0075-95 show endospore appendages of two morphological types

Negative stain EM imaging of *B. cereus* strain NVH 0075-95 showed typical endospores with a dense core of ∼1 μm diameter, tightly wrapped by an exosporium layer that on TEM images emanates as a flat 2-3 μm long saclike structure from the endospore body (Figure 1A). The endospores showed an abundance of micrometer-long appendages (Enas) (Figure 1A). The average endospore counted 20 - 30 Enas ranging from 200 nm to 6 μm in length (Figure 1E), with a median length of approximately 600 nm. The density of Enas appeared highest at the pole of the spore body that lies near the exosporium. There, Enas seem to emerge from the exosporium as individual fibers or as a bundle of individual fibers that separates a few tens of nanometers above the endospore surface (Figures 1B and S1B). Closer inspection revealed that the Enas showed two distinct morphologies (Figure 1 C, D). The main or “Staggered-type” (S-type) morphology represents approximately 90% of the observed fibers. S-type Enas have a width of ∼110 Å and give a polar, staggered appearance in negative stain 2D classes, with alternating scales pointing down to the spore surface. At the distal end, S-type Enas terminate in multiple filamentous extensions or “ruffles” of 50 - 100 nm in length and ∼35 Å thick (Figure 1C). The minor or “Ladder-like” (L-type) Ena morphology is thinner, ∼80 Å in width, and terminates in a single filamentous extension with dimensions similar to ruffles seen in S-type fibers (Figure 1D). L-type Enas lack the scaled, staggered appearance of the S-type Enas, instead showing a ladder of stacked disk-like units of ∼40 Å height. Whereas S-type Enas can be seen to traverse the exosporium and connect to the spore body, L-type Enas appear to emerge from the exosporium (Figure S1A). Both Ena morphologies co-exist on individual endospores (Figure S1C). Neither Ena morphology is reminiscent of sortase-mediated or type IV pili previously observed in Gram-positive bacteria (Mandlik et al., 2008; Melville and Craig, 2013). In an attempt to identify their composition, shear force extracted and purified Enas were subjected to trypsin digestion for identification by mass spectrometry. However, despite the good enrichment of both S- and L-type Enas, no unambiguous candidates for Ena were identified amongst the tryptic peptides, which largely contained contaminating mother cell proteins, EA1 S-layer and spore coat proteins. Attempts to resolve the Ena monomers by SDS-PAGE were unsuccessful, including strong reducing conditions (up to 200 mM β-mercaptoethanol), heat treatment (100 °C), limited acid hydrolysis (1h 1M HCl), or incubation with chaotropes such as 8M urea or 6M guanidinium chloride. Ena fibers also retained their structural properties upon autoclaving, desiccation or treatment with proteinase K (Figure S1C).

**Figure 1.**
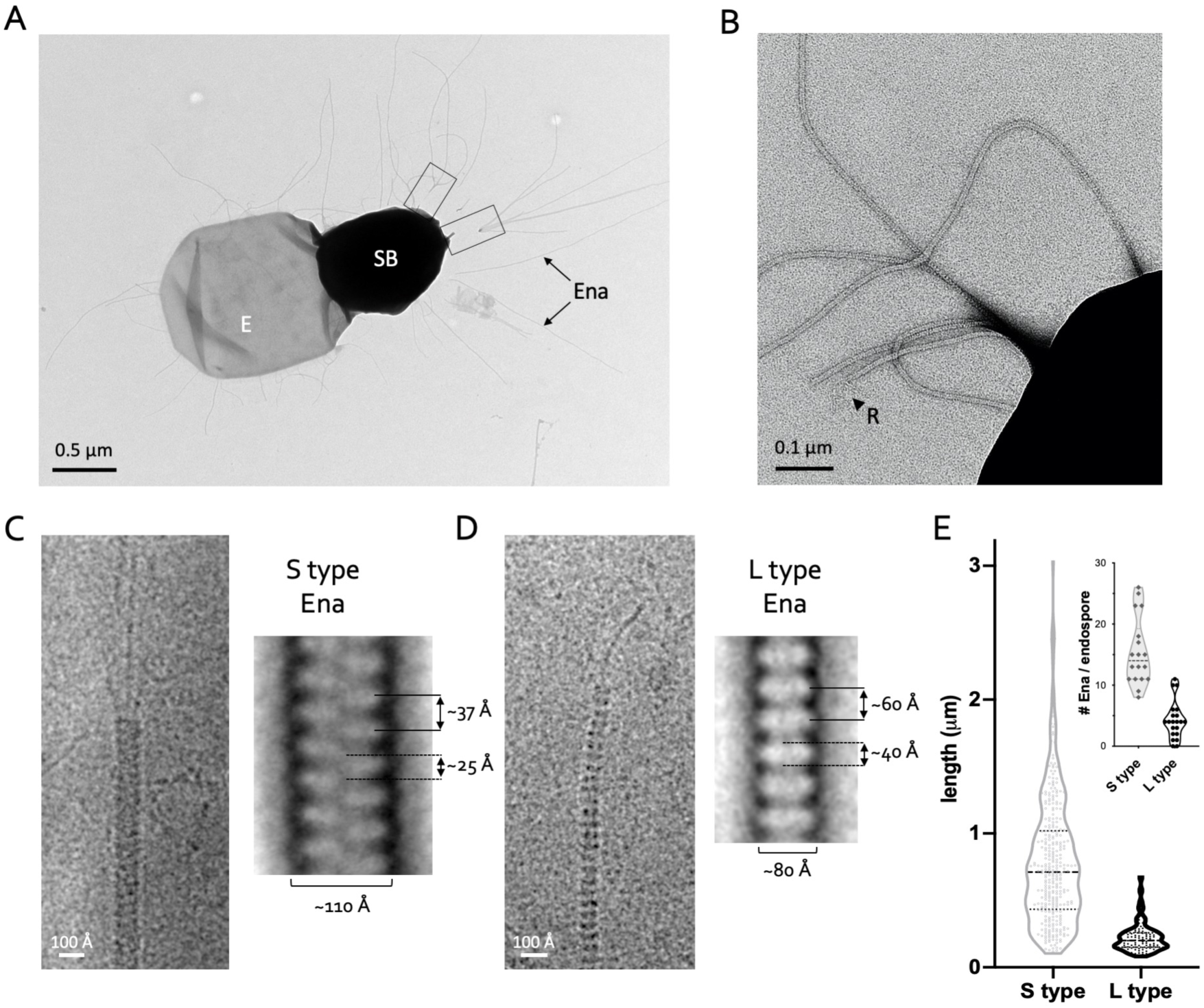
*B. cereus* endospores carry S and L-type Enas. **(A, B)** negative stain TEM image of *B. cereus* NVH 0075-95 endospore, showing spore body (SB), exosporium (E), and endospore appendages (Ena), which emerge from the endospore individually or as fiber clusters (boxed). At the distal end, Enas terminate in a single or multiple thin ruffles (R). **(C, D)** Single fiber cryoTEM images and negative stain 2D class averages of S-type (C) and L-type Enas (D). **(E)** Length distribution of S- and L-type Enas and number of Enas per endospore (inset), (*n*=1023, from 150 endospores, from 5 batches). See also Figure S1.

### Cryo-EM of endospore appendages identifies their molecular identity

To further study the nature of the Enas, fibers purified from *B. cereus* NVH 0075-95 endospores were imaged by cryogenic electron microscopy (cryo-EM) and analyzed using 3D reconstruction. Isolated fibers showed a 9.4:1 ratio of S- and L-type Enas, similar to what was seen on endospores. Boxes with a dimension of 300 × 300 pixels (246 × 246 Å^2^) were extracted along the length of the fibers, with an inter-box overlap of 21 Å, and subjected to 2D classification using RELION 3.0 (Zivanov et al., 2018). Power spectra of the 2D class averages revealed a well-ordered helical symmetry for S-type Enas (Figure 2A, B), whereas L-type Enas primarily showed translational symmetry (Figure 1D). Based on a helix radius of approximately 54.5 Å, we estimated layer lines Z’ and Z” in the power spectrum of S-type Enas to have a Bessel order of -11 and 1, respectively (Figure 2A, B). In the 2D classes holding the majority of extracted boxes the Bessel order 1 layer line was found at a distance of 0.02673 Å-1 from the equator, corresponding to a pitch of 37.4 Å, in good agreement with spacing of the apparent ‘lobes’ seen also by negative stain (Figures 1C, 2B and S1). The correct helical parameters were derived by an empirical approach in which a systematic series of starting values for subunit rise and twist were used for 3D reconstruction and real space Bayesian refinement using RELION 3.0 (He and Scheres, 2017). Based on the estimated Fourier – Bessel indexing, input rise and twist were varied in the range of 3.05 – 3.65 Å and 29 – 35 degrees, respectively, with a sampling resolution of 0.1 Å and 1 degree between tested start values. This approach converged on a unique set of helical parameters that resulted in 3D maps with clear secondary structure and identifiable densities for subunit side chains (Figure 2C). The reconstructed map corresponds to a left-handed 1-start helix with a rise and twist of 3.22937 Å and 31.0338 degrees per subunit, corresponding to a helix with 11.6 units per turn (Figure 2D). After refinement and postprocessing in RELION 3.0, the map was found to be of resolution 3.2 Å according to the FSC_0.143_ criterion.

**Figure 2.**
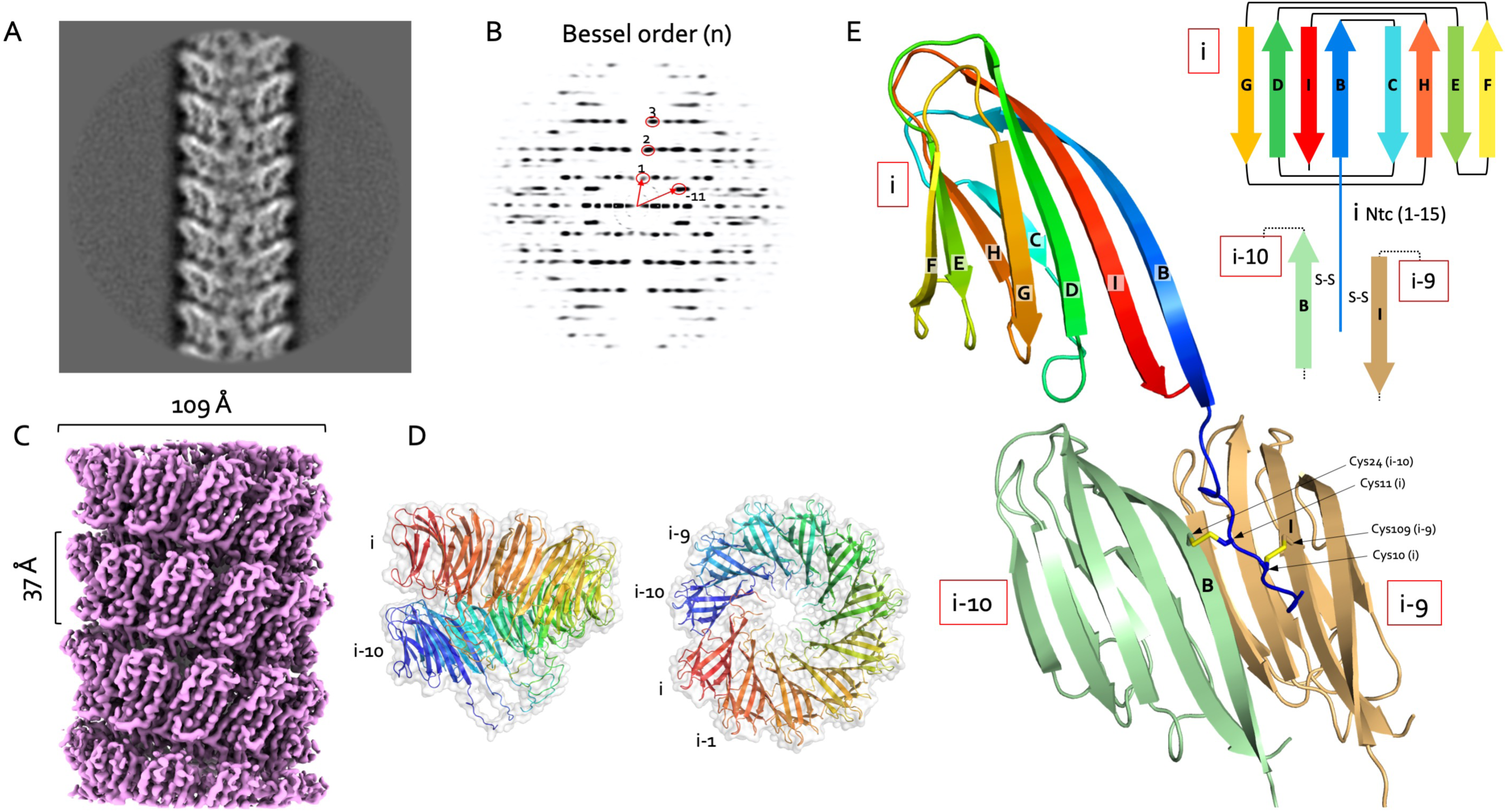
CryoTEM structure of S-type Enas. **(A, B)** Representative 2D class average (A) and corresponding power spectrum **(B)** of *B. cereus* NVH 0075-95 S-type Enas viewed by cryoTEM. Bessel orders used to derive helical symmetry are indicated. **(C)** Reconstituted cryoEM electron potential map of *ex vivo* S-type Ena (3.2 Å resolution). **(D)** Side and top view of a single helical turn of the *de novo*-built 3D model of S-type Ena shown in ribbon representation and molecular surface. Ena subunits are labelled i to i-10. **(F)** Ribbon representation and topology diagram of the S-type Ena1B subunit (blue to red rainbow from N- to C-terminus), and its interaction with subunits i-9 (sand) and i-10 (green) through disulfide crosslinking.

The resulting map showed well defined subunits comprising an 8-stranded β-sandwich domain of approximately 100 residues (Figure 2E). The side chain density was of sufficient quality to manually deduce a short motif with the sequence F-C-M-V/T-I-R-Y (Figure S2A). A search of the *B. cereus* NVH 0075-95 proteome (GCA_001044825.1) identified two hypothetical proteins of unknown function, namely KMP91697.1 and KMP91698.1, encoded by TU63_02435 and TU63_02440 respectively (Figure S2B). Further inspection of the electron potential map and manual model building of the Ena subunit showed this to fit well with the protein KMP91698.1. TU_63_02440 is located 15 bp downstream of the TU63_02435 locus. Both genes encode hypothetical proteins of similar size (117 and 126 amino acids and estimated molecular weights of 12 and 14 kDa, for KMP91698.1 and KMP91697.1, respectively), with 39% pairwise amino acid sequence identity, a shared domain of unknown function (DUF) 3992 and similar Cys patterns (Figure S2B). Further downstream of TU_63_02440, on the minus strand, the locus TU63_0245 encodes a third DUF3992 containing hypothetical protein (KMP91699.1), of 160 amino acids and an estimated molecular weight of 17 kDa. As such, KMP91697.1, KMP91698.1 and KMP91699.1 are regarded as candidate Ena subunits, hereafter dubbed Ena1A, Ena1B and Ena1C, respectively (Figure S2B,C).

### Ena1B self-assembles into endospore appendage-like nanofibers *in vitro*

To confirm the subunit identity of the endospore appendages isolated from *B. cereus* NVH0075-95, we cloned a synthetic gene fragment corresponding to the coding sequence of Ena1B and an N-terminal TEV protease cleavable 6xHis-tag into a vector for recombinant expression in the cytoplasm of *E. coli*. The recombinant protein was found to form inclusion bodies, which were solubilized in 8M urea before affinity purification. Removal of the chaotropic agent by rapid dilution resulted in the formation of abundant soluble crescent-shaped oligomers reminiscent of a partial helical turn seen in the isolated S-type Enas (Figure S2A-E), suggesting the refolded recombinant Ena1B (*rec*Ena1B) adopts the native subunit-subunit β-augmentation contacts (Figure S2E). We reasoned that *rec*Ena1B self-assemble into helical appendages arrested at the level of a single turn due to steric hindrance by the 6xHis-tag at the subunits N-terminus. Indeed, proteolytic removal of the affinity tag readily resulted in the formation of fibers of 110 Å diameter and with helical parameters similar to S-type Enas, though lacking the distal ruffles seen in *ex vivo* fibers (Figure S2F). CryoEM data collection and 3D helical reconstruction was performed to assess whether *in vitro rec*Ena1B nanofibers were isomorphous with *ex vivo* S-type Enas. Real space refinement of helical parameters using RELION 3.0 converged on a subunit rise and twist of 3.43721 Å and 32.3504 degrees, respectively, approximately 0.2 Å and 1.3 degrees higher than found in *ex vivo* S-type Enas, and corresponding to a left-handed helix with a pitch of 38.3 Å and 11.1 subunits per turn. Apart from the minor differences in helical parameters the 3D reconstruction map of *in vitro* Ena1B fibers (estimated resolution of 3.2 Å; Figure S3A, B) was near isomorphous to *ex vivo* S-type Enas in terms of size and connectivity of the fiber subunits (Figure S3D). Closer inspection of the 3D cryoEM maps for *rec*Ena1B and *ex vivo* S-type Ena showed an improved side chain fit for Ena1B residues in the former (Figure S3B, C, D) and revealed regions in the *ex vivo* Ena maps that showed partial side-chain character of Ena1A, particularly in loop L1, L3, L5 and L7 (Figure S2B, S3B,C). Although the Ena1B character of the *ex vivo* maps is dominant, this suggested that *ex vivo* S-type Enas consist of a mixed population of Ena1A and Ena1B fibers, or that S-type Enas have a mixed composition comprising both Ena1A and Ena1B. Immunogold labelling using sera generated with *rec*Ena1A or *rec*Ena1B showed subunits-specific labeling within single Enas, confirming these have a mixed composition of Ena1A and Ena1B (Figure S3E). No staining of S-type Enas was seen with Ena1C serum (Figure S3E). No systematic patterning or molar ratio for Ena1A and Ena1B could be discerned from immunogold labelling or helical reconstructions with an asymmetric unit containing more than one subunit, suggesting the distribution of Ena1A and Ena1B in the fibers to be random. Apart from some side chain densities with mixed Ena1A and Ena1B character, the cryoEM electron potential maps of the *ex vivo* Enas showed a unique main chain conformation, indicating the Ena1A and Ena1B have near isomorphous folds.

### Enas represent a novel family of Gram-positive pili

Upon recognizing that native S-type Enas show a mixed Ena1A and Ena1B composition, we continued with 3D cryoEM reconstruction of *rec*Ena1B for model building. The Ena subunit consists of a typical jellyroll fold (Richardson, 1981) comprised of two juxtaposed β-sheets consisting of strands BIDG and CHEF (Figure 2F). The jellyroll domain is preceded by a flexible 15 residue N-terminal extension hereafter referred to as N-terminal connector (‘Ntc’). Subunits align side by side through a staggered β-sheet augmentation (Remaut and Waksman, 2006), where the sheet composed of strands BIDG of a subunit i is augmented with strands CHEF of the preceding subunit i-1, and strands CHEF of subunit i are augmented with strands BIDG of the next subunit in row i+1 (Figure 2F, Figure S4A, B). As such, the packing in the endospore appendages can be regarded as a slanted β-propeller of 8-stranded β-sheets, with 11.6 blades per helical turn and an axial rise of 3.2 Å per subunit (Figure 2E). Subunit-subunit contacts in the β-propeller are further stabilized by two complementary electrostatic patches on the Ena subunits (Figure S4C). In addition to these lateral contacts, subunits across helical turns are also connected through the Ntc’s. The Ntc of each subunit i makes disulfide bond contacts with subunits i-9 and i-10 in the preceding helical turn (Figure 2F, Figure S4B). These contacts are made through disulfide bonding of Cys 10 and Cys 11 in subunit i, with Cys 109 and Cys 24 in the strands I and B of subunits i-9 and i-10, respectively (Figure 2F, S4B). Thus, disulfide bonding via the Ntc results in a longitudinal stabilization of fibers by bridging the helical turns, as well as in a further lateral stabilization in the β-propellers by covalent cross-linking of adjacent subunits. The Ntc contacts lie on the luminal side of the helix, leaving a central void of approximately 1.2 nm diameter (Figure S4D). Residues 12-17 form a flexible spacer region between the Ena jellyroll domain and the Ntc. Strikingly, this spacer region creates a 4.5 Å longitudinal gap between the Ena subunits, which are not in direct contact other than through the Ntc (Figure 3C, S2B). The flexibility in the Ntc spacer and the lack of direct longitudinal protein-protein contact of subunits across the helical turns create a large flexibility and elasticity in the Ena fibers (Figure 3). 2D class averages of endospore-associated fibers show longitudinal stretching, with a change in pitch of up to 8 Å (range: 37.1 – 44.9 Å; Figure 3D), and an axial rocking of up to 10 degrees per helical turn (Figure 3A, B).

**Figure 3.**
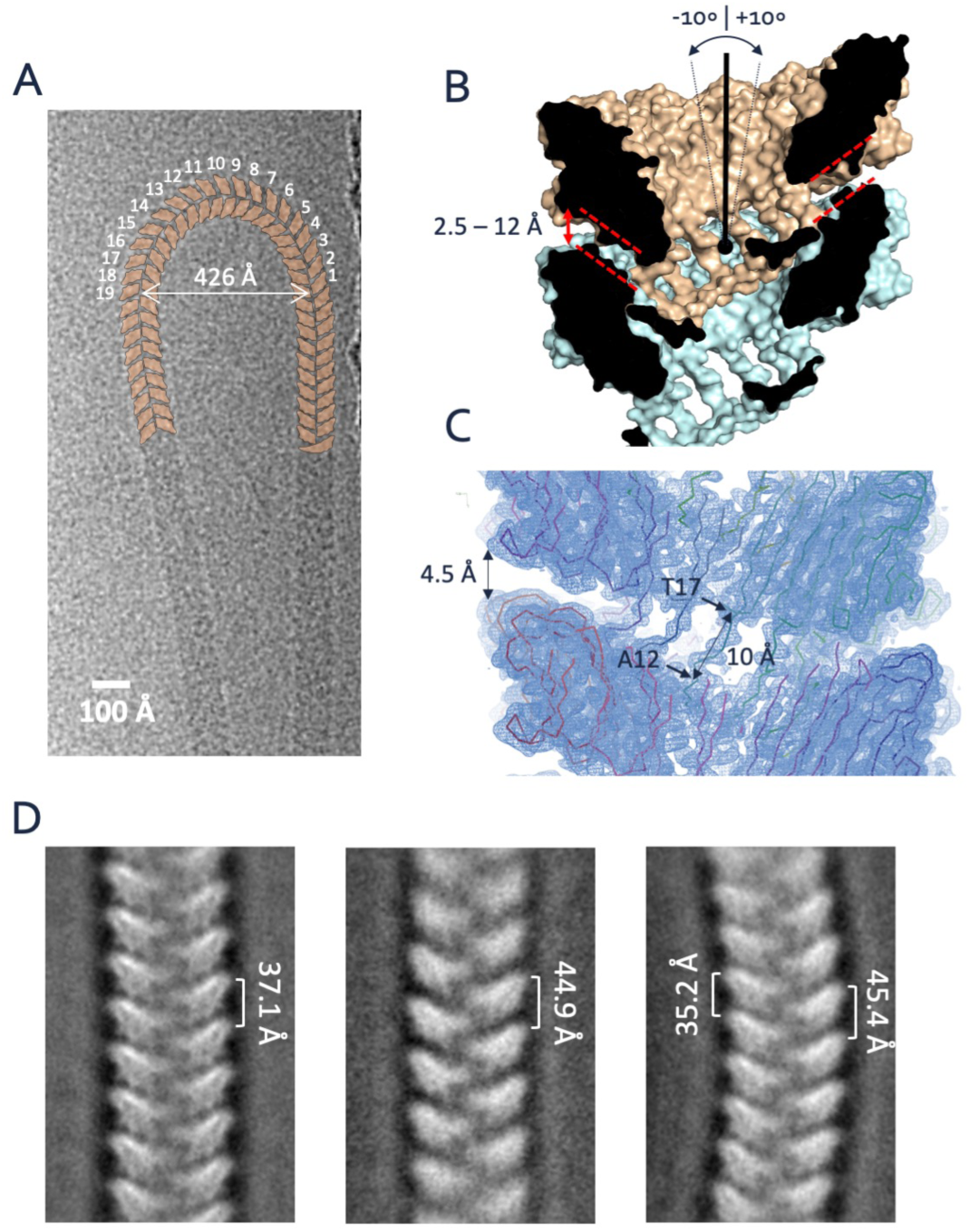
Ntc linkers give high flexibility and elasticity to S-type Enas. **(A)** CryoTEM image of an isolated S-type Ena making a U-turn comprising just 19 helical turns (shown schematically in orange). **(B, C)** Cross-section and 3D cryoTEM electron potential map of the S-type Ena model, highlighting the longitudinal spacing between Ena1B jellyroll domains as a result of the Ntc linker (residues 12-17). **(D)** Negative stain 2D class averages of endospore-associated S-type Enas show variation in pitch and axial curvature.

Thus, *B. cereus* endospore appendages represent a novel class of bacterial pili, comprising a left-handed single start helix with non-covalent lateral subunit contacts formed by β-sheet augmentation, and covalent longitudinal contacts between helical turns by disulfide bonded N-terminal connecter peptides, resulting in an architecture that combines extreme chemical stability (Figure S1) with high fiber flexibility.

### The *ena1* coding region for S-type Enas

In *B. cereus* NVH 0075-95 Ena1A, Ena1B and Ena1C are encoded in a genomic region flanked upstream by *dedA* (genbank protein-id: KMP91696.1) and a gene encoding a 93-residue protein of unknown function (DUF1232, genbank: KMP91695.1) (Figure 4A). Downstream, the *ena*-gene cluster is flanked by a gene encoding an acid phosphatase (TU63_02450). Within the *ena*-gene cluster, *ena1A* and *ena1B* are found in forward, and *ena1C* in reverse orientation, respectively (Figure 4A). PCR analysis of NVH 0075-95 cDNA made from mRNA isolated after 4 and 16 h of culture, representative for vegetative growth and sporulating cells, respectively, indicated that *ena1A* and *ena1B* are co-expressed from a bicistronic transcript during sporulation but not during vegetative growth (Figure 4B). A weak amplification signal was observed in vegetative cells when the forward primer was located in *dedA* upstream of *ena1A* and the reverse primer was located within *ena1B* (Figure 4B, lane 2) suggesting that some *enaA* and *enaB* is coexpressed with *dedA*. This was observed in vegetative cells or very early in sporulation but not during later sporulation stages and may represent a fraction of improperly terminated *dedA* mRNA. Reverse transcription quantitative PCR (RT-qPCR) analysis showed increased expression of *ena1A, ena1B* and *ena1C* in sporulating cells compared to vegetative cells (Figure 4B). CryoEM maps and immuno-gold TEM analysis of *ex vivo* S-type Enas indicated these contain both Ena1A and Ena1B (Figure S3B-D). To determine the relative contribution of Ena1 subunits to *B. cereus* Enas we made individual chromosomal knockouts of *ena1A, ena1B*, as well as *ena1C* in strain NVH 0075-95 and investigated their respective endospores by TEM. All *ena1* mutants made endospores of similar dimensions to WT and with intact exosporium (Figure 5A, Figure S5). Both the *ena1A* and *ena1B* mutant resulted in endospores completely lacking S-type Enas, in agreement with the mixed content of *ex vivo* fibers. The *ena1C* mutant also resulted in the loss of S-type Ena on the endospores (Figure 5A), even though staining with anti-Ena1C serum did not identify the presence of the protein inside S-type Enas (Figure S3D). All three mutants still showed the presence of L-type Enas, of similar size and number density as WT endospores, although statistical analysis does not rule out L-type Enas to have a slight increase in length in the *ena1B* and *ena1C* mutants (length p=0.003 and <0.0001, resp.) (Figure 5B). Thus, Ena1A, Ena1B and Ena1C are mutually required for *in vivo* S-type Ena assembly, but not for L-type Ena assembly. Complementation of the *ena1B* mutant with a low copy plasmid (pMAD-I-*Sce*I) containing *ena1A-ena1B* restored S-type Ena expression. Plasmid-based expression of these subunits resulted in an average ∼2-fold increase in the number of S-type Enas per spore, and a drastic increase in Ena length, now reaching several microns (Figure 5A, B, Figures S5D). Thus, the number and length of S-type Enas depend on the concentration of available Ena1A and Ena1B subunits. Notably, several endospores overexpressing Ena1A and Ena1B appeared to lack an exosporium or showed the entrapment of S-type Enas inside the exosporium (Figure S5C, D). This demonstrates that S-type Enas emanate from the spore body, and that a disbalance in the concentration or timing of *ena* expression can result in mis-assembly and/or mislocalization of endospore surface structures. Contrary to S-type Enas, close inspection of the WT and mutant endospores suggests that L-type Enas emanate from the surface of the exosporium rather than the spore body. The molecular identity of the L-type Ena, or the single or multiple terminal ruffles seen, respectively, in L- and S-type Enas was not determined in the present study.

**Figure 4.**
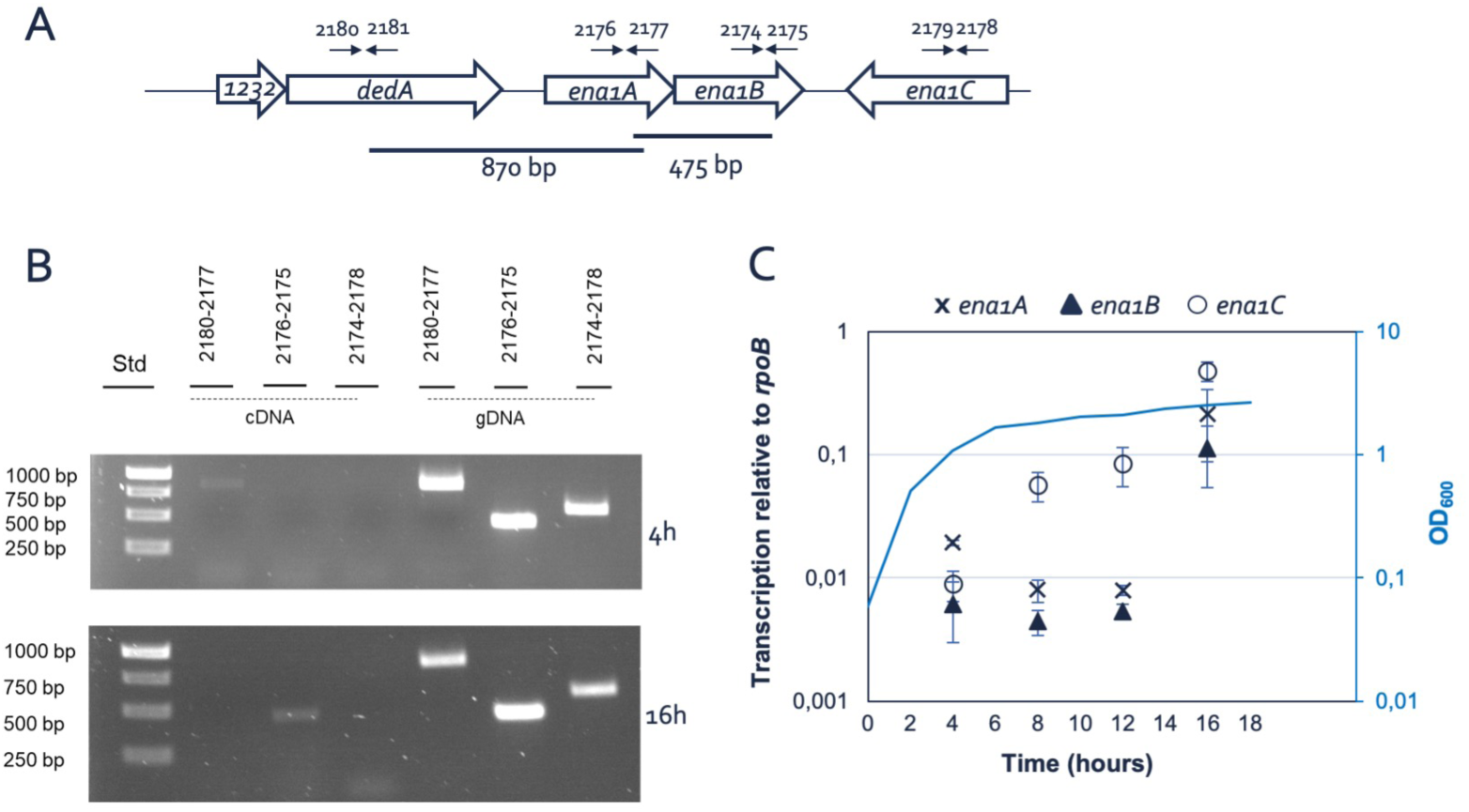
*ena* is bicistronic and expressed during sporulation. **(A)** Chromosomal organization of the *ena* genes and primers used for transcript analysis (arrows). **(B)** Agarose gel electrophoresis (1%) analysis of PCR products using indicated primer pairs and cDNA made of mRNA isolated from NVH 0075-95 after 8 and 16 hours growth in liquid cultures or genomic DNA as control. Of note, the transcription of *ena*1C was surprisingly higher than *ena1A* and *ena1B*, which are components of the major appendages. **(C)** Transcription level of *ena1A* (x), *ena1B* (▲), *ena1C* (⍰) and *dedA* (⍰) relative to *rpoB* determined by RT-qPCR during 16⍰hours of growth of *B. cereus* strain NVH 0075-95. The dotted line represents the bacterial growth measured by increase in OD_600_. Whiskers represent standard deviation of three independent experiments.

**Figure 5.**
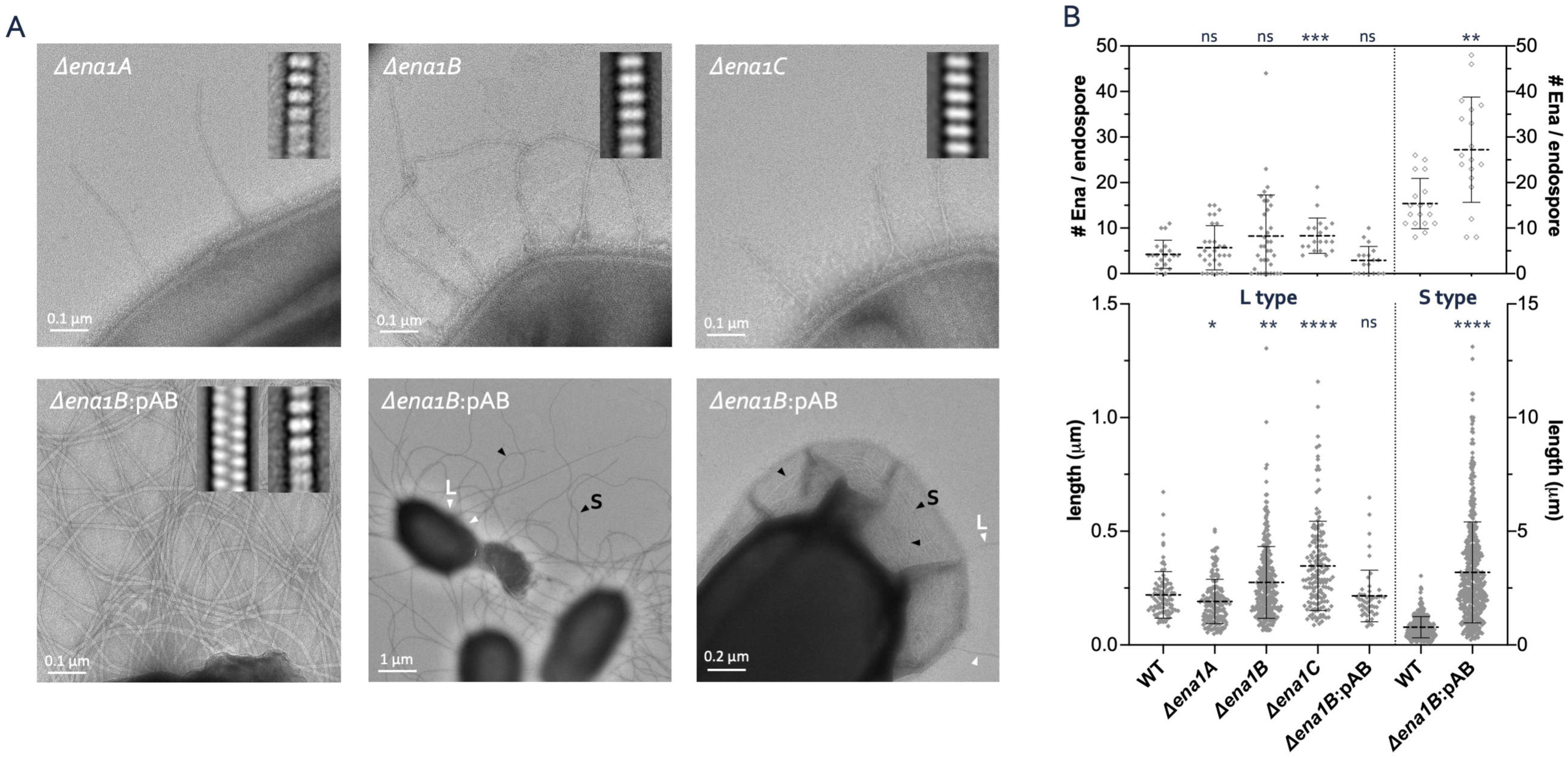
Composition of S- and L-type Ena. **(A)** Representative negative stain images of endospores of NVH 0075-95 mutants lacking *ena1A, ena1B, ena1A* and **B** or *ena1C*, as well as the *ena1B* mutant complemented with *ena1A-ena1B* from plasmid (pAB). Inset are 2D class averages of Enas observed on the respective mutants. **(B)** Length distribution and number of Enas found on WT and mutant NVH 0075-95 endospores. Statistics: pair-wise Mann-Whitney U tests against WT (*n*: ≥18 spores; *n*: ≥50 Enas; ns: not significant, * p<0.05, ** p<0.01, *** p<0.001 and **** p<0.0001. mean ± s.d.)

### Phylogenetic distribution of the *ena1A-C* genes

To investigate the occurrence of *ena1A-C* within the *B. cereus s*.*l*. group and other relevant species of the genus *Bacillus*, pairwise tBLASTn searches for homologs of Ena1A-C were performed on a database containing all available closed, curated *Bacillus* spp. genomes, with the addition of scaffolds for species for which closed genomes were lacking (*n*=735, Table S3). Homologs and orthologs with high coverage (>90%) and high amino acid sequence identity (>80%) to Ena1A or Ena1B of *B. cereus* NVH 0075-95 were found in 48 strains including 11 of 85 *B. cereus* strains, 13 of 119 *B. wiedmannii* strains, 14 of 14 *B. cytotoxicus* strains, one of one *B. luti* (100%) strain, three of six *B. mobilis* strains, three of 33 *B. mycoides* strains, one of one *B. tropics* strain and both *B. paranthracis* strains analyzed. Of these strains, only 31 also carried a gene encoding a homolog with high sequence identity and coverage to Ena1C of *B. cereus* NVH 0075-95 (Figure 6). All investigated *B. cytotoxicus* genomes (14/14) encoded hypothetical Ena1A and Ena1B proteins, but only 12/14 encoded an Ena1C ortholog, which showed only a moderate amino acid conservation compared to the Ena1C of *B. cereus* NVH 0075-95 (mean 63.9% amino acid sequence identity) (Figure 6, Figure S5).

**Figure 6.**
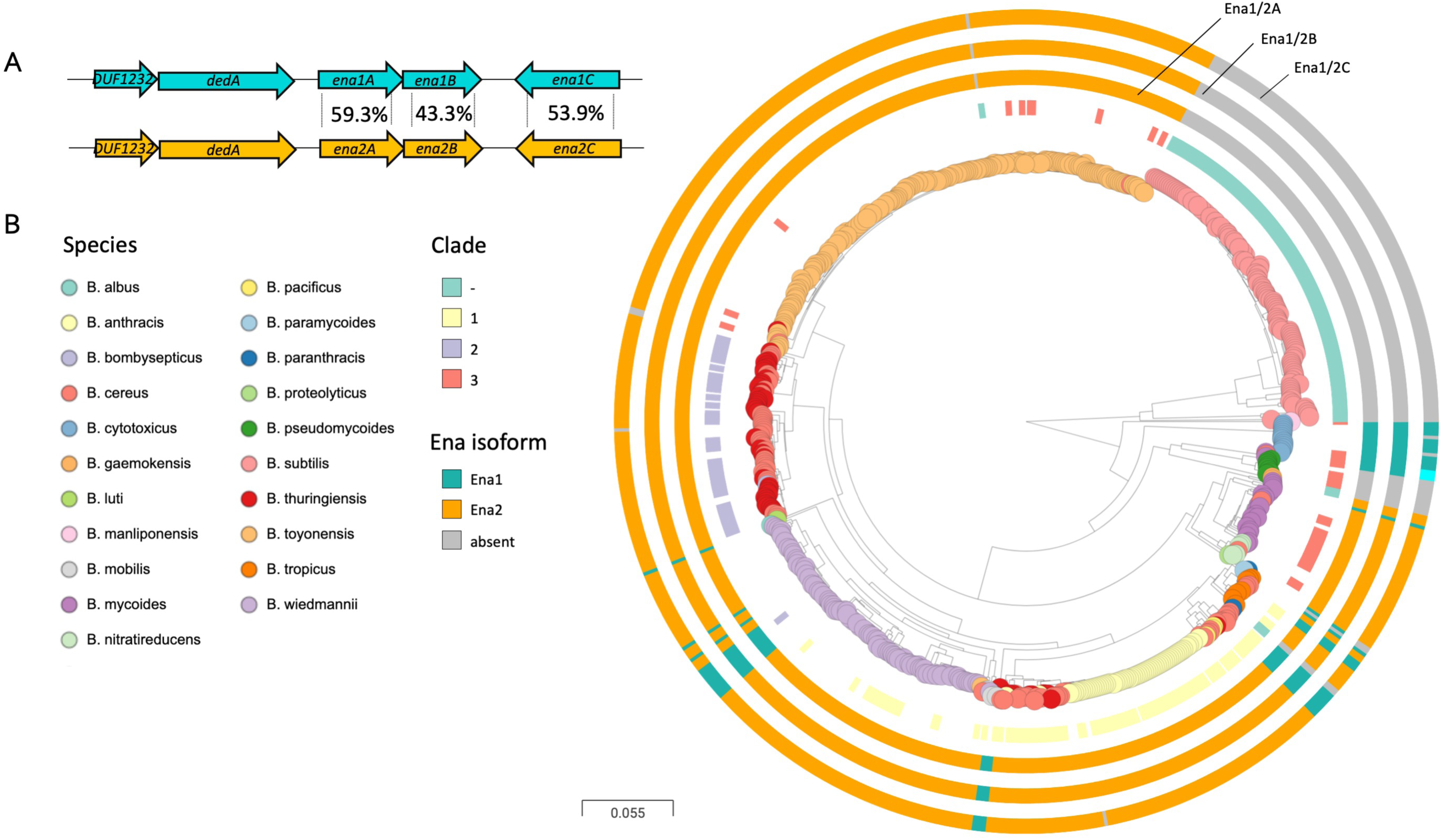
Ena is widespread in pathogenic Bacilli. **(A)** Ena1 and Ena2 loci with average amino acid sequence identity indicated between the population of EnaA-C ortho- and homologs. Ena1C shows considerably more variation and is in *B. cytotoxicus* different from both Ena1C and Ena2C (see Figure S5C), while other genomes have *enaC* present at different loci (applies to two isolates of *B. mycoides)*. **(B)** Distribution of *ena1/2A-C* among *Bacillus* species. Whole genome clustering of the *B. cereus s*.*l*. group and *B. subtilis* created by Mashtree (Katz et al., 2019; Ondov et al., 2016) and visualized in Microreact (Argimon et al., 2016). Rooted on *B. subtilis*. Traits for species (colored nodes), Bazinet clades and presence of *ena* are indicated on surrounding four rings in the following order from inner to outer: clade according to Bazinet 2017 (when available) (Bazinet, 2017) (see legend), and presence of *enaA, enaB* and *enaC* (for all three, *ena1*: teal, *ena2*: orange, different locus: cyan). When no homo- or ortholog was found, the ring is grey. Ena1A-C and Ena2A-C are defined as present when a homologous protein of the corresponding genome has high coverage (>90%) and >80% and 50-65% sequence identity, respectively, with Ena1A-C of the NMH 0095/75 strain (see Table S4). Interactive tree accessible at https://microreact.org/project/vn2oWw7zM3cwejEFNoRGWA/0024f86c

Upon searching for Ena1A-C homologs in *B. cereus* group genomes, a candidate orthologous gene cluster encoding hypothetical EnaA-C proteins was discovered. These three proteins had, respectively, an average of 59.3±0.9%, 43.3±1.6% and 53.9±2.2% amino acid sequence identity with Ena1A, Ena1B and Ena1C of *B. cereus* NVH0075-95, and shared gene synteny (Figure 6b). The orthologous *ena* gene cluster was named *ena2A-C*. Except for *B. subtilis (n=127)* and *B. pseudomycoides (n=8)*, all genomes analyzed (*n=735*) carried either *ena1* (*n=48*) or the *ena2* (*n=476*) gene cluster. *Ena1A-C* or the *ena2A-C* were never present simultaneously and no chimeric *ena1A-C*/*2A-C* clusters were discovered among the genomes analyzed (Figure 6). In addition to the main split between Ena1A-C and Ena2A-C in the protein trees, distinct sub-clusters were seen among Ena1A, Ena1B and, especially, Ena1C sequences (Figure S5). The Ena1A sequences separated into two main sub-clusters: one present in the majority of *B. cytotoxicus* strains and another found in *B. wiedmanni* and *B. cereus* strains (Figure S5A). More variation was evident for EnaB proteins: Ena1B sequences formed two clusters; one containing *B. cereus* and *B. wiedmannii* isolates, and the other with *B. cytotoxicus* (Figure S5B). Also, a separate sub-cluster of Ena2B proteins was seen (Figure S5B), containing isolates of *B. mycoides, B. cereus, B. thuringiensis, B. pacificus*, and *B. wiedmannii* that shared around ∼78% and ∼48% sequence identity with the remainder of Ena2B and Ena1B, respectively. EnaC was the most variable of the three proteins: Ena1C formed a monophyletic clade containing isolates of *B. wiedmanni, B. cereus, B. anthracis, B. paranthracis, B. mobilis, B. tropicus*, and *B. luti*, but had considerable sequence variation in species and strains carrying *Ena2AB* as well as in subset of strains carrying *Ena1AB*.

The *ena2A-C* homo- or orthologs were much more common among *B. cereus* group strains than the *ena1A-C* genes; all investigated *B. toyonensis* (*n=204*), *B. albus* (*n=1*), *B. bombysepticus* (*n=1), B. nitratireducens* (*n=6*), *B. thuringiensis* (*n=50*) genomes and in the majority of *B. cereus* (87%, 74/85), *B. wiedmannii* (105/119, 89.3%), *B. tropicus* (71%, 5/7,) and *B. mycoides* (91%, 30/33) had the Ena2A-C form of the protein (Figure 6). No *ena* orthologs were found in *B. subtilis* (*n=127*) or *B. pseudomycoides* (n=8) genomes or in any other genomes outside the *B. cereus* group except for three misclassified *Streptococcus pneumoniae* genomes (GCA_001161325, GCA_001170885, GCA_001338635) and one misclassified *B. subtilis* genome (GCA_004328845). These genomes and the *B. subtilis* were re-classified as *B. cereus* when re-analyzed with three different methods for t*axonomic* classification (Masthree, 7-lociMLST and Kraken, see Methods). The genomes of a few *Peanibacillus* spp. strains had genes encoding hypothetical proteins with a low level of amino acid sequence similarity to Ena1A-C, and genes encoding hypothetical proteins with some similarity to Ena1A and Ena1B were also found in the genome of a *Cohnella abietis* strain (GCF_004295585.1). These hits outside of *Bacillus* genus was in the DUF3992 domain of these genes, which is found in *Anaeromicrobium, Cochnella*, and of the order Bacillales.

A few genomes had deviations in the *ena-*gene clusters compared to other strains of their species. Two of three *B. mycoides* strains (GCF_007673655 and GCF_007677835.1) lacked the *ena1C* allele downstream of the *ena1A-B* operon (data not shown). However, potential *ena1c* orthologs encoding hypothetical proteins with 50% identity to Ena1C of *B. cereus* NVH 0075-95 were found elsewhere in their genomes. One genome annotated as *B. cereus* (strain Rock3-44 Assembly: GCA_000161255.1) grouped with these strains of *B. mycoides* (Figure 6) and shared their *ena1A-C* distribution pattern with. *B. thuringiensis* usually carries *ena2* gene, but a genome annotated as *B. thuringiensis* (strain LM1212, GCF_003546665) lacked all *ena* genes. This strain was nearly identical to the reference strain of *B. tropicu*s, which also lacked both the *ena* gene clusters. One *B. toyonensis* strain AFS086269 GCF_002568845 also lacked all three genes, while the remainder of the 204 strains of *B. toyonensis* all had *ena2A-C*.

## Discussion

Endospores formed by *Bacillus* and *Clostridium* species frequently carry surface-attached ribbon- or pilus-like appendages (Driks, 2007), the role of which has remained largely enigmatic due to the lack of molecular annotation of the pathways involved in their assembly. Half a century following their first observation (Hachisuka and Kuno, 1976; Hodgikiss, 1971), we employ high resolution *de novo* structure determination by cryoEM to structurally and genetically characterize the appendages found on *B. cereus* spores. We found that *B. cereus* Enas come in two main morphologies: 1) staggered or S-type Enas that are several micrometers long and emerge from the spore body and traverses the exosporium, and 2) smaller, less abundant ladder- or L-type Enas that appears to directly emerge from the exosporium surface. Our phylogenetic analyses of S-type fibers reveal Ena subunits belonging to a conserved family of proteins encompassing the domain of unknown function DUF3992.

Covalent bonding, and the highly compact jellyroll fold result in a high chemical and physical stability of the Ena fibers, withstanding desiccation, high temperature treatment, and exposure to proteases. The formation of linear filaments of multiple hundreds of subunits requires stable, long-lived subunit-subunit interactions with high flexibility to avoid that a dissociation of subunit-subunit complexes results in pilus breakage. This high stability and flexibility are likely to be adaptations to the extreme conditions that can be met by endospores in the environment or during the infectious cycle. Two molecular pathways are known to form surface fibers or “pili” in Gram-positive bacteria: 1) sortase-mediated pilus assembly, which encompasses the covalent linkage of pilus subunits by means of a transpeptidation reaction catalyzed by sortases (Ton-That and Schneewind, 2004), and 2) Type IV pilus assembly, encompassing the non-covalent assembly of subunits through a coiled-coil interaction of a hydrophobic N-terminal helix (Melville and Craig, 2013). Sortase-mediated pili and Type IV pili are formed on vegetative cells, however, and to date, no evidence is available to suggest that these pathways are also responsible for the assembly of endospore appendages.

Until the present study, the only species for which the genetic identity and protein composition of spore appendages has been known, is the non-toxigenic environmental species *Clostridium taeniosporum*, which carry large (4.5 µm long, 0.5 µm wide and 30 nm thick) ribbon-like appendages, which are structurally distinct from those found in most other *Clostridium* and *Bacillus* species. *C. taeniosporum* lacks the exosporium layer and the appendages seem to be attached to another layer, of unknown composition, outside the coat (Walker et al., 2007). The *C. taeniosporum* endospore appendages consist of four major components, three of which have no known homologs in other species and an orthologs of the *B. subtilis* spore membrane protein SpoVM (Walker et al., 2007). The appendages on the surface of *C. taeniosporum* endospores, therefore, represent distinct type of fibers than those found on the surface of spores of species belonging to the *B. cereus* group.

Our structural studies uncover a novel class of pili, where subunits are organized into helically wound fibers, held together by lateral β-sheet augmentation inside the helical turns, and longitudinal disulfide cross-linking across helical turns. Covalent cross-linking in pilus assembly is known for sortase-mediated isopeptide bond formation seen in Gram-positive pili (Ton-That and Schneewind, 2004). In Enas, the cross-linking occurs through disulfide bonding of a conserved Cys-Cys motif in the N-terminal connector of a subunit i, to two single Cys residues in the core domain of the Ena subunits located at position i-9 and i-10 in the helical structure. As such, the N-terminal connectors form a covalent bridge across helical turns, as well as a branching interaction with two adjacent subunits in the preceding helical turn (i.e. i-9 and i-10). The use of N-terminal connectors or extensions is also seen in chaperone-usher pili and *Bacteroides* Type V pili, but these system employ a non-covalent fold complementation mechanism to attain long-lived subunit-subunit contacts, and lack a covalent stabilization (Sauer et al., 1999; Xu et al., 2016). Because in Ena the N-terminal connectors are attached to the Ena core domain via a flexible linker, the helical turns in Ena fibers have a large pivoting freedom and ability to undergo longitudinal stretching. These interactions result in highly chemically stable fibers, yet with a large degree of flexibility. Whether the stretchiness and flexibility of Enas carry a functional importance remains unclear. Of note, in several chaperone-usher pili, a reversible spring-like stretching provided by helical unwinding and rewinding of the pili has been found important to withstand shear and pulling stresses exerted on adherent bacteria (Fallman et al., 2005; Miller et al., 2006). Possibly, the longitudinal stretching seen in Ena may serve a similar role.

Typical Ena filaments have, to the best of our knowledge, never been observed on the surface of vegetative *B. cereus* cells indicating that they are endospore-specific structures. In support of that assumption, RT-qPCR analysis NVH 0075-95 demonstrated increased *ena1A-C* transcript during sporulation, compared to vegetative cells. A transcriptional analysis of *B. thuringiensis* serovar *chinensis* CT-43 at 7 h, 9 h, 13 h (30 % of cells undergoing sporulation) and 22 hours of growth has previously been performed (Wang et al., 2013). It is difficult to directly compare expression levels of *ena1A, B* and *C* in *B. cereus* NVH 0075-95 with the expression level of *ena2A-C* in *B. thuringiensis* serovar *chinensis CT-43* (CT43_CH0783-785) since the expression of the latter strain was normalized by converting the number of reads per gene into RPKM (Reads Per Kilo bases per Million reads) and analyzed by DEGseq software package, while the present study determines the expression level of the *ena* genes relative to the house keeping gene *rpoB*. However, both studies indicate that *enaA* and *enaB* are only transcribed during sporulation. By searching a separate set of published transcriptomic profiling data we found that *ena2A-C* also are expressed in *B. anthracis* during sporulation (Bergman et al., 2006), although Enas have not previously been reported from *B. anthracis* spores.

Without knowledge on the function of Enas, we can only speculate about their biological role. The Enas of *B. cereus* group species resemble pili, which in Gram-negative and Gram-positive vegetative bacteria play roles in adherence to living surfaces (including other bacteria) and non-living surfaces, twitching motility, biofilm formation, DNA uptake (natural competence) and exchange (conjugation), secretion of exoproteins, electron transfer (Geobacter) and bacteriophage susceptibility (Lukaszczyk et al., 2019; Proft and Baker, 2009). Some bacteria express multiple types of pili that perform different functions. The most common function of pili-fibers is adherence to a diverse range of surfaces from metal, glass, plastics rocks to tissues of plants, animals or humans. In pathogenic bacteria, pili often play a pivotal role in colonization of host tissues and function as important virulence determinants. Similarly, it has been shown that appendages, expressed on the surface of *C. sporogenes* endospores, facilitate their attachment to cultured fibroblast cells (Panessa-Warren et al., 2007). The Enas are, however, not likely to be involved in active motility or uptake/transport of DNA or proteins as they are energy demanding processes that are not likely to occur in the endospore’s metabolically dormant state. Enas appear to be a widespread feature among spores of strains belonging to the *B. cereus* group (Figure 6), a group of closely related *Bacillus* species with a strong pathogenic potential (Ehling-Schulz et al., 2019). For most *B. cereus* group species, the ingestion, inhalation or the contamination of wounds with endospores forms a primary route of infection and disease onset. Enas cover much of the cell surface so that they can be reasonably expected to form an important contact region with the endospore environment and may play a role in the dissemination and virulence of *B. cereus* species. Our phylogenetic analysis shows a widespread occurrence of Enas in pathogenic *Bacilli*, and a striking absence in non-pathogenic species such as *B. subtilis*, a soil-dwelling species and gastrointestinal commensal that has functioned as the primary model system for studying endospores. Ankolekar *et al*., showed that all of 47 food isolates of *B. cereus* produced endospores with appendages (Ankolekar and Labbe, 2010). Appendages were also found on spores of ten out of twelve food-borne, enterotoxigenic isolates of *B. thuringiensis*, which is closely related to *B. cereus*, and best known for its insecticidal activity (Ankolekar and Labbe, 2010).

The cryo-EM images of *ex vivo* fibers showed 2-3 nm wide fibers (ruffles) at the terminus of S- and L-type Enas. The ruffles resemble tip fibrilla of P-pili and type 1 seen in many Gram-negatives bacteria of the family Enterobacteriaceae (Proft and Baker, 2009). In Gram-negative pilus filaments, the tip fibrilla provides adhesion proteins with a flexible location to enhance the interaction with receptors on mucosal surfaces (Mulvey et al., 1998). No ruffles were observed on the *in vitro* assembled fibers suggesting that their formation require additional components than the Ena1B subunits.

We present the molecular identification of a novel class of spore-associated appendages or pili widespread in pathogenic *Bacilli*. Future molecular and infection studies will need to determine if and how Enas play a role in the virulence of spore-borne pathogenic *Bacilli*. The advances in uncovering the genetic identity and the structural aspects of the Enas presented in this work now enable *in vitro* and *in vivo* molecular studies to tease out their biological role(s), and to gain insights into the basis for Ena heterogeneity amongst different *Bacillus* species.

## Acknowledgements

We thank Markus Fislage and Adam Schrofel at the VIB-VUB Facility for Bio Electron Cryogenic Microscopy (BECM) for assistance in data collection and Jan Haug Anonsen at NORCE research, Norway / Department of Biosciences, University of Oslo for assistance with sample analysis. We are grateful to Ute Krengel (UiO) for the mentorship of J.L. and feedback on the manuscript. This works was funded by VIB, EOS Excellence in Research Program by FWO through grant G0G0818N to HR, the NMBUs talent development program to MA and travel grants from The national graduate school in infection biology and antimicrobials (IBA) through NFR grant 249069 to J.L.

## Author contributions

B.P and M.S. performed TEM imaging, structural studies, and recombinant ena1B production and analysis. J.L. and T.L. produced endospores, performed TEM imaging, isolated Enas and conducted genetic studies. A-K.L. and O.B. conducted the phylogenetic analysis. H.R. and M.A. designed and supervised experiments, and wrote the paper, with contributions from all authors.

## METHODS

### Culture of *B. cereus* and appendages extraction

For extraction of Enas the *B. cereus* strain NVH 0075-95 was plated on blood agar plates and incubated at 37 °C for 3 months. Upon maturation, the spores were resuspended and washed in milli-Q water three times (centrifugation 2,400 *g* at 4 °C). To get rid of various organic and inorganic debris, the pellet was then resuspended in 20% Nycodenz (Axis-Shield) and subjected to Nycodenz density gradient centrifugation where the gradient was composed of a mixture of 45% and 47% (w/v) Nycodenz in 1:1 v/v ratio. The pellet consisting only of the spore cells was then washed with 1 M NaCl and TE buffer (50 mM Tris-HCl; 0.5 mM EDTA) containing 0.1% SDS respectively. To detach the appendages, the washed spores were sonicated at 20k Hz ± 50 Hz and 50 watts (Vibra Cell VC50T; Sonic & Materials Inc.; U.S.) for 30 seconds on ice followed by centrifugation at 4500 *g* and appendages were collected in the supernatant. To further get rid of the residual components of spore and vegetative mother cells n-hexane was added and vigorously mixed with the supernatant in 1:2 v/v ratio. The mixture was then left to settle to allow phase separation of water and hexane. The hexane fraction containing the appendages was then collected and kept at 55 °C under pressured air for 1.5 hours to evaporate the hexane. The appendages were finally resuspended in mill-Q water for further cryo-EM sample preparation.

### Recombinant expression, purification and *in vitro* assembly of Ena1B appendages

A synthetic open reading frame encoding Ena1B was codon optimized for recombinant expression in *E. coli*, synthesized and cloned into pET28a expression vector at Twist biosciences (Table S2). The insert was designed to have a N-terminal 6X histidine tag on Ena1B along with a TEV protease cleavage site (ENLYFQG) in between. Large scale recombinant expression was carried out in the T7 Express lysY/Iq *E. coli* strain from NEB. A single colony was inoculated into 20 mL of LB and grown at 37 °C with shaking at 150 rpm overnight for primary culture. Next morning 6 L of LB was inoculated with 20 mL/L of primary culture and grown at 37 °C with shaking until the OD_600_ reached 0.8 after which protein expression was induced with 1 mM isopropyl β-D-1-thiogalactopyranoside (IPTG). The culture was incubated for a further 3 hours at 37 °C and harvested by centrifugation at 5,000 rpm. The whole-cell pellet was resuspended in lysis buffer (20 mM potassium phosphate, 500 mM NaCl, 10 mM β -mercaptoethanol, 20 mM imidazole, pH 7.5) and sonicated on ice for lysis. The lysate was centrifuged to separate the soluble and insoluble fractions by centrifugation at 18,000 rpm for 45 min in a JA-20 rotor from Beckman Coulter. The pellet was further dissolved in denaturing lysis buffer consisting 8M urea in lysis buffer. The dissolved pellet was then passed over a HisTrap HP column (GE Healthcare) and equilibrated with denaturing lysis buffer. The bound protein was eluted from the column with elution buffer (20 mM potassium phosphate, pH 7.5, 8 M urea, 250 mM imidazole) in a gradient mode (20-250 mM imidazole) at room temperature.

For *in vitro* Ena1B assembly, purified His-Ena1B in denaturing conditions was first dialyzed against a buffer containing 20 mM Hepes, pH 7.0, 50 mM NaCl overnight at 4 °C. To facilitate Ena1B self-assembly into S-type Ena filaments the 6xHis-tag was cleaved off by TEV protease. TEV protease along with 100 mM β-mercaptoethanol was then added in equimolar ratio and incubated for 2 hours at 37 °C. Removal of the 6xHis-tag led to the assembly of the *rec*Ena1B into long Ena-like filaments Figure S2F.

### Ena treatment experiments to test its robustness

*Ex vivo* Enas extracted from *B. cereus* strain NVH 0075-95 (see above) were resuspended in deionized water, autoclaved at 121 °C for 20 minutes to ensure inactivation of residual bacteria or spores, and subjected to treatment with buffer or as indicated below and shown in Figure S1. To determine Ena integrity upon the various treatments, samples were imaged using negative stain TEM and Enas were boxed and subjected to 2D classification as described below. To test protease resistance, *ex vivo* Ena were subjected to 1 mg/mL Ready-to-use Proteinase K digestion (Thermo Scientific) for 4 hours at 37 °C and imaged by TEM. To study the effects of desiccation on the appendages, *ex vivo* Ena were vacuum dried at 43 °C using Savant DNA120 Speedvac Concentrator (Thermo scientific) run for 2 hours at a speed of 2k RPM.

### Negative-Stain Transmission Electron Microscopy (TEM)

For visualization of spores and recombinantly expressed appendages by negative stain TEM, formvar/carbon coated copper grids with 400-hole mesh (Electron Microscopy Sciences) were glow discharged (ELMO; Agar Scientific) with a plasma current of 4 mA at vacuum for 45 seconds. 3 µL of sample was applied on the grids and allowed to bind to the support film for 1 minute after which the excess liquid was blotted away with Whatman grade 1 filter paper. The grids were then washed three times using three 15 µL drops of milli-Q followed by blotting of excess liquid. The washed grids were held in 15 µL drops of 2% uranyl acetate three times for, respectively, 10 seconds, 2 seconds and 1 minute durations, with a blotting step in between each dip. Finally, the uranyl acetate coated grids were blotted until dry. The grids were then imaged using a 120 kV JEOL 1400 microscope equipped with LaB6 filament and TVIPS F416 CCD camera. 2D classes of the appendages were generated in RELION 3.0 (Zivanov et al., 2018) as described below.

### Preparation of cryo-TEM grids and cryo-EM data collection

QUANTIFOIL® holey Cu 400 mesh grids with 2 µm holes and 1 µm spacing were first glow discharged in vacuum using plasma current of 5 mA for 1 minute (ELMO; Agar Scientific). 3 µL of 0.6 mg /mL graphene oxide (GO) solution was applied onto the grid and incubated 1 minute for absorption at room temperature. Extra GO was then blotted out and left for drying using a Whatman grade 1 filter paper. For cryo-plunging, 3 µL of protein sample was applied on the GO coated grids at 100% humidity and room temperature in a Gatan CP3 cryo-plunger. After 1 minute of absorption it was machine-blotted with Whatman grade 2 filter paper for 5 seconds from both sides and plunge frozen into liquid ethane at 180 °K. Grids were then stored in liquid nitrogen until data collection. Two datasets were collected for *ex vivo* and *rec*Ena1B appendages with slight changes in the collection parameters. High resolution cryo-EM 2D micrograph movies were recorded on a JEOL Cryoarm300 microscope automated with energy filter and a K2 or K3 direct electron detector run in counting mode. For the *ex vivo* Ena, the microscope was equipped with a K2 summit detector and had the following settings: 300 keV, 100 mm aperture, 30 frames / image, 62.5 e^-^/Å^2^, 2.315 seconds exposure, and 0.82 Å/pxl. For the *rec*Ena1B dataset was recorded on a K3 detector, at a pixel size of 0.782 Å/pxl, and an exposure of 64.66 e-/Å2 taken over 61 frames / image.

### Image processing

MOTIONCORR2 (Zheng et al., 2017) implemented in RELION 3.0 (Zivanov et al., 2018) was used to correct for beam-induced image motion and averaged 2D micrographs were generated. The motion-corrected micrographs were used to estimate the CTF parameters using CTFFIND4.2 (Rohou and Grigorieff, 2015) integrated in RELION 3.0. Subsequent processing used RELION 3.0. and SPRING (Desfosses et al., 2014). For both the datasets, the coordinates of the appendages were boxed manually using *e2helixboxer* from the EMAN2 package (Tang et al., 2007). Special care was taken to select micrographs with good ice and straight stretches of Ena filaments. The filaments were segmented into overlapping single-particle boxes of dimension 300 × 300 pxl with an inter-box distance of 21 Å. For the *ex vivo* Enas a total of 53,501 helical fragments was extracted from 580 micrographs with an average of 2 - 3 long filaments per micrograph. For the *rec*Ena1B filaments, 100,495 helical fragments were extracted from 3,000 micrographs with an average of 4 - 5 filaments per micrograph. To filter out bad particles multiples rounds of 2D classification were run in RELION 3.0. After several rounds of filtering, a dataset of 42,822 and 65,466 good particles of the *ex vivo* and *rec*Ena1B appendages were selected, respectively.

After running ∼50 iterations of 2D classification well-resolved 2D class averages could be obtained. *segclassexam* of the SPRING package (Desfosses et al., 2014) was used to generate B-factor enhanced power spectrum of the 2D class averages. The generated power spectrum had an amplified signal-to-noise ratio with well resolved layer lines (Figure 2B). To estimate crude helical parameters, coordinates and phases of the peaks in the layer lines were measured using the *segclasslayer* option in SPRING. Based on the measured distances and phases possible sets of Bessel orders were deduced, after which the calculated helical parameters were used in a helical reconstruction procedure in RELION (He and Scheres, 2017). A featureless cylinder of 110 Å diameter generated using *relion_helix_toolbox* was used as an initial model for 3D classification. Input rise and twist deduced from Fourier – Bessel indexing were varied in the range of 3.05 – 3.65 Å and 29 – 35 degrees, respectively, with a sampling resolution of 0.1 Å and 1 degree between tested start values. So doing, several rounds of 3D classification were run until electron potential maps with good connectivity and recognizable secondary structure were obtained. The output translational information from the 3D classification was used to re-extract particles and 3D refinement was done taking a 25 Å low pass filtered map generated from the 3D classification run. To improve the resolution of the EM maps multiple rounds of 3D refinement were run. To further improve the resolution Bayesian polishing was performed in RELION. Finally, a solvent mask covering the central 50% of the helix *z*-axis was generated in *maskcreate* and used for postprocessing and calculating the solvent-flattened Fourier shell correlation (FSC) curve in RELION. After two rounds of polishing, maps of 3.2 Å resolution according to the FSC_0.143_ gold-standard criterion as well as local resolution calculated in RELION were obtained (Figure S3A).

### Model building

Prior to model building unfiltered maps for recEna1B calculated by Relion were masked down to three helical turns and used for cryo-EM density modification as implemented in ResolveCryoEM (Terwilliger et al., 2019) from the PHENIX package (Afonine et al., 2018), resulting in a map of 3.05 Å final resolution (FSC_0.143_ criterium) for *rec*Ena1B. At first the primary skeleton for a single asymmetric subunit from the density modified map was generated in Coot (Emsley et al., 2010). Primary sequence of Ena1B was manually threaded onto the asymmetric unit and fitted into the map taking into consideration the chemical properties of the residues. The SSM superpose tool in coot was used to place the additional protomers of the S-type Ena from a single subunit. The built model was then subjected to multiple rounds real space structural refinement in PHENIX, each residue was manually inspected after every round of refinement. Model validation was done in Molprobity (Davis et al., 2007) implemented in Phenix. All the visualizations and images for figures were generated in ChimeraX (Goddard et al., 2018), Chimera (Pettersen et al., 2004), or Pymol.

### Immuno-labelling of the Enas

For antibody generation, *rec*Ena1A and *rec*Ena1C were cloned, expressed and purified according to the method described above for *rec*Ena1B. Aliquots of purified *rec*Ena1A, *rec*Ena1B and *rec*Ena1C were concentrated to 1 mg/mL in PBS for rabbit immunization (Davids Biotechnologie GmbH). For immunostaining EM imaging, 3 µL aliquots of purified *ex vivo* Enas were deposited on Formvr/Carbon grids (400 Mesh, Cu; Electron Microscopy Sciences), washed with 20 μL 1x PBS, and incubated for 1 hour with 0.5% (w/v) BSA in 1x PBS. After additional washing with 1x PBS, separate grids were individually incubated for 2 hours at 37 °C with 1:1000 dilutions in PBS of anti-Ena1A, anti-Ena1B or anti-Ena1C sera, respectively. Following washing with 1x PBS, grids were incubated for 1 hour 37 °C with a 1:2000 dilution of a 10 nm gold-labelled goat anti-rabbit IgG (G7277; Sigma Aldrich), washed with 1x PBS, and negative stained and imaged on a 120 kV JEOL 1400 microscope as described above.

### Quantitative RT-PCR

RT-qPCR experiments were performed on isolated mRNA from *B. cereus* cultures harvested from three independent Bacto media cultures (37 °C, 150 rpm) at 4, 8, 12 and 16hours post-inoculation. RNA extraction, cDNA synthesis and RT-qPCR analysis was performed essentially as described before (Madslien et al., 2014), with the following changes: pre-heated (65 °C) TRIzol Reagent (Invitrogen) and bead beating 4 times for 2 minutes in a Mini-BeadBeater-8 (BioSpec) with cooling on ice in between.

Each RT-qPCR of the RNA samples was performed in triplicate, no template was added in negative controls, and *rpoB* was used as internal control. Slopes of the standard curves and PCR efficiency (E) for each primer pair were estimated by amplifying serial dilutions of the cDNA template. For quantification of mRNA transcript levels, Ct (threshold cycle) values of the target genes and the internal control gene (*rpoB*) derived from the same sample in each RT-qPCR reaction were first transformed using the term E^-Ct^. The expression levels of target genes were then normalized by dividing their transformed Ct-values by the corresponding values obtained for the internal control gene (Duodu et al., 2010; Madslien et al., 2014; Pfaffl, 2001). The amplification was conducted by using StepOne PCR software V.2.0 (Applied Biosystems) with the following conditions: 50 °C for 2 minutes, 95 °C for 2 minutes, 40 cycles of 15 seconds at 95 °C, 1 minute at 60 °C and 15 seconds at 95 °C. All primers used for RT-qPCR analyses are listed in Table S2.

Regular PCR reactions were performed on cDNA to confirm that *ena1A* and *ena1B* were expressed as an operon using the primers 2180/2177 and 2176/2175 and DreamTaq DNA polymerase (Thermo Fisher) amplified in an Eppendorf Mastercycler using the following program: 95 °C for 2 minutes, 30 cycles of 95 °C for 30 seconds, 54 °C for 30 seconds, and 72 °C for 1 minute.

### Construction of deletion mutants

The *B. cereus* strain NVH 0075-95 was used as background for gene deletion mutants. The e*na1B* gene was deleted in-frame by replacing the reading frames with ATGTAA (51–31) using a markerless gene replacement method (Janes and Stibitz, 2006) with minor modifications. To create the deletion mutants the regions upstream (primer A and B, Table S2) and downstream (primer C and D, Table S2) of the target *ena* genes were amplified by PCR. To allow assembly of the PCR fragments, primers B and C contained complementary overlapping sequences. An additional PCR step was then performed, using the upstream and downstream PCR fragments as template and the A and D primer pair (Table S2). All PCR reactions were conducted using an Eppendorf Mastercycler gradient and high fidelity AccuPrime Taq DNA Polymerase (ThermoFisher Scientific) according to the manufacturer’s instructions. The final amplicons were cloned into the thermosensitive shuttle vector pMAD (Arnaud et al., 2004) containing an additional I-SceI site as previously described (Lindback et al., 2012). The pMAD-I-*Sce*I plasmid constructs were passed through One Shot™ INV110 *E. coli* (ThermoFisher Scientific) to achieve unmethylated DNA to enhance the transformation efficiency in *B. cereus*. The unmethylated plasmid were introduced into *B. cereus* NVH 0075-95 by electroporation (Mahillon et al., 1989). After verification of transformants by PCR, the plasmid pBKJ233 (unmethylated), containing the gene for the I-*Sce*I enzyme, was introduced into the transformant strains by electroporation. The I-*Sce*I enzyme makes a double-stranded DNA break in the chromosomally integrated plasmid. Subsequently, homologous recombination events lead to excision of the integrated plasmid resulting in the desired genetic replacement. The gene deletions were verified by PCR amplification using primers A and D (Table S2) and DNA sequencing (Eurofins Genomics).

### Search for orthologs and homologs of Ena1

Publicly available genomes of species belonging to the Bacillus s.l. group (Table S3 and S4) was downloaded from NCBI RefSeq database (*n=735*, NCB (https://www.ncbi.nlm.nih.gov/refseq/, <u>Table S1</u>). Except for strains of particular interest due to phenotypic characteristics (GCA_000171035.2_ASM17103v2, GCA_002952815.1_ASM295281v1, GCF_000290995.1_Baci_cere_AND1407_G13175) and species of which closed genomes were nonexistent or very scarce (*n=158*), all assemblies included were closed and publicly available genomes from the curated database of NCBI RefSeq database. Assemblies were quality checked using QUAST (Gurevich et al., 2013), and only genomes of correct size (∼4.9-6Mb) and a GC content of ∼35% were included in the downstream analysis. Pairwise tBLASTn searches were performed (e-value 1e-10, max_hspr 1, default settings) to search for homo- and orthologs of the following query-protein sequences from strain NVH 0075-95: Ena1A, Ena1B, Ena1C (Table S3). The Ena1B protein sequences used as query originated from an inhouse amplicon sequenced product, while the Ena1A and Ena1C protein sequence queries originated from the assembly for strain NVH 0075-95 (Accession number GCF_001044825.1, protein KMP91698.1 and KMP91699.1, Table S5). We considered proteins orthologs or homologs when a subject protein matched the query protein with high coverage (>70%) and moderate sequence identity (>30%).

### Comparative genomics of the ena-genes and proteins

Phylogenetic trees of the aligned Ena1A-C proteins were constructed using approximately maximum likelihood by FastTree (Price et al., 2010) (default settings) for all hits resulting from the tBLASTn search. The amino acid sequences were aligned using mafft v.7.310 (Katoh et al., 2019), and approximately-maximum-likelihood phylogenetic trees of protein alignments were made using FastTree, using the JTT+CAT model (Price et al., 2010). All Trees were visualized in Microreact (Argimon et al., 2016) and the metadata of species, and presence and absence for Ena1A-C and Ena2A-C overlaid the figures.

## References

Afonine, P.V., Poon, B.K., Read, R.J., Sobolev, O.V., Terwilliger, T.C., Urzhumtsev, A., and Adams, P.D. (2018). Real-space refinement in PHENIX for cryo-EM and crystallography. Acta Crystallogr D Struct Biol 74, 531–544.

Ankolekar, C., and Labbe, R.G. (2010). Physical characteristics of spores of food-associated isolates of the Bacillus cereus group. Appl Environ Microbiol 76, 982–984.

Argimon, S., Abudahab, K., Goater, R.J.E., Fedosejev, A., Bhai, J., Glasner, C., Feil, E.J., Holden, M.T.G., Yeats, C.A., Grundmann, H., et al. (2016). Microreact: visualizing and sharing data for genomic epidemiology and phylogeography. Microb Genom 2, e000093.

Arnaud, M., Chastanet, A., and Debarbouille, M. (2004). New vector for efficient allelic replacement in naturally nontransformable, low-GC-content, gram-positive bacteria. Appl Environ Microbiol 70, 6887–6891.

Atrih, A., and Foster, S.J. (1999). The role of peptidoglycan structure and structural dynamics during endospore dormancy and germination. Antonie Van Leeuwenhoek 75, 299–307.

Bazinet, A.L. (2017). Pan-genome and phylogeny of Bacillus cereus sensu lato. BMC Evol Biol 17, 176.

Bergman, N.H., Anderson, E.C., Swenson, E.E., Niemeyer, M.M., Miyoshi, A.D., and Hanna, P.C. (2006). Transcriptional profiling of the Bacillus anthracis life cycle in vitro and an implied model for regulation of spore formation. J Bacteriol 188, 6092–6100.

Davis, I.W., Leaver-Fay, A., Chen, V.B., Block, J.N., Kapral, G.J., Wang, X., Murray, L.W., Arendall, W.B., 3rd, Snoeyink, J., Richardson, J.S., et al. (2007). MolProbity: all-atom contacts and structure validation for proteins and nucleic acids. Nucleic Acids Res 35, W375–383.

Desfosses, A., Ciuffa, R., Gutsche, I., and Sachse, C. (2014). SPRING - an image processing package for single-particle based helical reconstruction from electron cryomicrographs. J Struct Biol 185, 15–26.

DesRosier, J.P., and Lara, J.C. (1981). Isolation and properties of pili from spores of Bacillus cereus. J Bacteriol 145, 613–619.

Driks, A. (2007). Surface appendages of bacterial spores. Mol Microbiol 63, 623–625.

Duodu, S., Holst-Jensen, A., Skjerdal, T., Cappelier, J.M., Pilet, M.F., and Loncarevic, S. (2010). Influence of storage temperature on gene expression and virulence potential of Listeria monocytogenes strains grown in a salmon matrix. Food Microbiol 27, 795–801.

Ehling-Schulz, M., Lereclus, D., and Koehler, T.M. (2019). The Bacillus cereus Group: Bacillus Species with Pathogenic Potential. Microbiol Spectr 7.

Emsley, P., Lohkamp, B., Scott, W.G., and Cowtan, K. (2010). Features and development of Coot. Acta crystallographica Section D, Biological crystallography 66, 486–501.

Fallman, E., Schedin, S., Jass, J., Uhlin, B.E., and Axner, O. (2005). The unfolding of the P pili quaternary structure by stretching is reversible, not plastic. EMBO Rep 6, 52–56.

Gerhardt, P., and Ribi, E. (1964). Ultrastructure of the Exosporium Enveloping Spores of Bacillus Cereus. Journal of bacteriology 88, 1774–1789.

Goddard, T.D., Huang, C.C., Meng, E.C., Pettersen, E.F., Couch, G.S., Morris, J.H., and Ferrin, T.E. (2018). UCSF ChimeraX: Meeting modern challenges in visualization and analysis. Protein Sci 27, 14–25.

Gurevich, A., Saveliev, V., Vyahhi, N., and Tesler, G. (2013). QUAST: quality assessment tool for genome assemblies. Bioinformatics 29, 1072–1075.

Hachisuka, Y., and Kuno, T. (1976). Filamentous appendages of Bacillus cereus spores. Jpn J Microbiol 20, 555–558.

He, S., and Scheres, S.H.W. (2017). Helical reconstruction in RELION. J Struct Biol 198, 163–176.

Hodgikiss, W. (1971). Filamentous appendages on the spores and exosporium of certain Bacillus species. In Spore research, A.N. Barker, G.W. Gould, and J. Wolf, eds. (London and New York: Academic Press), pp. 211–218.

Janes, B.K., and Stibitz, S. (2006). Routine markerless gene replacement in Bacillus anthracis. Infect Immun 74, 1949–1953.

Katoh, K., Rozewicki, J., and Yamada, K.D. (2019). MAFFT online service: multiple sequence alignment, interactive sequence choice and visualization. Brief Bioinform 20, 1160–1166.

Katz, L.S., Griswold, T., Morrison, S.S., Caravas, J.A., Zhang, S., C., d.B.H., Deng, X., and Carleton, A. (2019). Mashtree: a rapid comparison of whole genome sequence files. Journal of Open Source Software 4.

Lindback, T., Mols, M., Basset, C., Granum, P.E., Kuipers, O.P., and Kovacs, A.T. (2012). CodY, a pleiotropic regulator, influences multicellular behaviour and efficient production of virulence factors in Bacillus cereus. Environ Microbiol 14, 2233–2246.

Lukaszczyk, M., Pradhan, B., and Remaut, H. (2019). The Biosynthesis and Structures of Bacterial Pili. Subcell Biochem 92, 369–413.

Madslien, E.H., Granum, P.E., Blatny, J.M., and Lindback, T. (2014). L-alanine-induced germination in Bacillus licheniformis -the impact of native gerA sequences. BMC Microbiol 14, 101.

Mahillon, J., Chungjatupornchai, W., Decock, J., Dierickx, S., Michiels, F., Peferoen, M., and Joos, H. (1989). Transformation of Bacillus thuringiensis by electroporation. FEMS Microbiology Letters 60, 205–210.

Mandlik, A., Swierczynski, A., Das, A., and Ton-That, H. (2008). Pili in Gram-positive bacteria: assembly, involvement in colonization and biofilm development. Trends Microbiol 16, 33–40.

Melville, S., and Craig, L. (2013). Type IV pili in Gram-positive bacteria. Microbiol Mol Biol Rev 77, 323–341.

Miller, E., Garcia, T., Hultgren, S., and Oberhauser, A.F. (2006). The mechanical properties of E. coli type 1 pili measured by atomic force microscopy techniques. Biophys J 91, 3848–3856.

Mulvey, M.A., Lopez-Boado, Y.S., Wilson, C.L., Roth, R., Parks, W.C., Heuser, J., and Hultgren, S.J. (1998). Induction and evasion of host defenses by type 1-piliated uropathogenic Escherichia coli. Science 282, 1494–1497.

Ondov, B.D., Treangen, T.J., Melsted, P., Mallonee, A.B., Bergman, N.H., Koren, S., and Phillippy, A.M. (2016). Mash: fast genome and metagenome distance estimation using MinHash. Genome Biol 17, 132.

Panessa-Warren, B.J., Tortora, G.T., and Warren, J.B. (2007). High resolution FESEM and TEM reveal bacterial spore attachment. Microsc Microanal 13, 251–266.

Pettersen, E.F., Goddard, T.D., Huang, C.C., Couch, G.S., Greenblatt, D.M., Meng, E.C., and Ferrin, T.E. (2004). UCSF Chimera--a visualization system for exploratory research and analysis. J Comput Chem 25, 1605–1612.

Pfaffl, M.W. (2001). A new mathematical model for relative quantification in real-time RT-PCR. Nucleic Acids Res 29, e45.

Price, M.N., Dehal, P.S., and Arkin, A.P. (2010). FastTree 2--approximately maximum-likelihood trees for large alignments. PLoS One 5, e9490.

Proft, T., and Baker, E.N. (2009). Pili in Gram-negative and Gram-positive bacteria - structure, assembly and their role in disease. Cell Mol Life Sci 66, 613–635.

Remaut, H., and Waksman, G. (2006). Protein-protein interaction through beta-strand addition. Trends Biochem Sci 31, 436–444.

Richardson, J.S. (1981). The anatomy and taxonomy of protein structure. Adv Protein Chem 34, 167–339.

Rode, L.J., Pope, L., Filip, C., and Smith, L.D. (1971). Spore appendages and taxonomy of Clostridium sordellii. Journal of bacteriology 108, 1384–1389.

Rohou, A., and Grigorieff, N. (2015). CTFFIND4: Fast and accurate defocus estimation from electron micrographs. J Struct Biol 192, 216–221.

Sauer, F.G., Futterer, K., Pinkner, J.S., Dodson, K.W., Hultgren, S.J., and Waksman, G. (1999). Structural basis of chaperone function and pilus biogenesis. Science 285, 1058–1061.

Setlow, P. (2014). Germination of spores of Bacillus species: what we know and do not know. Journal of bacteriology 196, 1297–1305.

Smirnova, T.A., Zubasheva, M.V., Shevliagina, N.V., Nikolaenko, M.A., and Azizbekian, R.R. (2013). [Electron microscopy of the surfaces of bacillary spores]. Mikrobiologiia 82, 698–706.

Stewart, G.C. (2015). The Exosporium Layer of Bacterial Spores: a Connection to the Environment and the Infected Host. Microbiol Mol Biol Rev 79, 437–457.

Tang, G., Peng, L., Baldwin, P.R., Mann, D.S., Jiang, W., Rees, I., and Ludtke, S.J. (2007). EMAN2: an extensible image processing suite for electron microscopy. J Struct Biol 157, 38–46.

Terwilliger, T.C., Ludtke, S.J., Read, R.J., Adams, P.D., and Afonine, P.V. (2019). Improvement of cryo-EM maps by density modification. bioRxiv.

Ton-That, H., and Schneewind, O. (2004). Assembly of pili in Gram-positive bacteria. Trends Microbiol 12, 228–234.

Walker, J.R., Gnanam, A.J., Blinkova, A.L., Hermandson, M.J., Karymov, M.A., Lyubchenko, Y.L., Graves, P.R., Haystead, T.A., and Linse, K.D. (2007). Clostridium taeniosporum spore ribbon-like appendage structure, composition and genes. Mol Microbiol 63, 629–643.

Wang, J., Mei, H., Zheng, C., Qian, H., Cui, C., Fu, Y., Su, J., Liu, Z., Yu, Z., and He, J. (2013). The metabolic regulation of sporulation and parasporal crystal formation in Bacillus thuringiensis revealed by transcriptomics and proteomics. Mol Cell Proteomics 12, 1363–1376.

Xu, Q., Shoji, M., Shibata, S., Naito, M., Sato, K., Elsliger, M.A., Grant, J.C., Axelrod, H.L., Chiu, H.J., Farr, C.L., et al. (2016). A Distinct Type of Pilus from the Human Microbiome. Cell 165, 690–703.

Zheng, S.Q., Palovcak, E., Armache, J.P., Verba, K.A., Cheng, Y., and Agard, D.A. (2017). MotionCor2: anisotropic correction of beam-induced motion for improved cryo-electron microscopy. Nat Methods 14, 331–332.

Zivanov, J., Nakane, T., Forsberg, B.O., Kimanius, D., Hagen, W.J., Lindahl, E., and Scheres, S.H. (2018). New tools for automated high-resolution cryo-EM structure determination in RELION-3. Elife 7.

