## Supplementary material for "*Bacillus* endospore appendages form a novel family of disulfide-linked pili"

**Supplemental information**

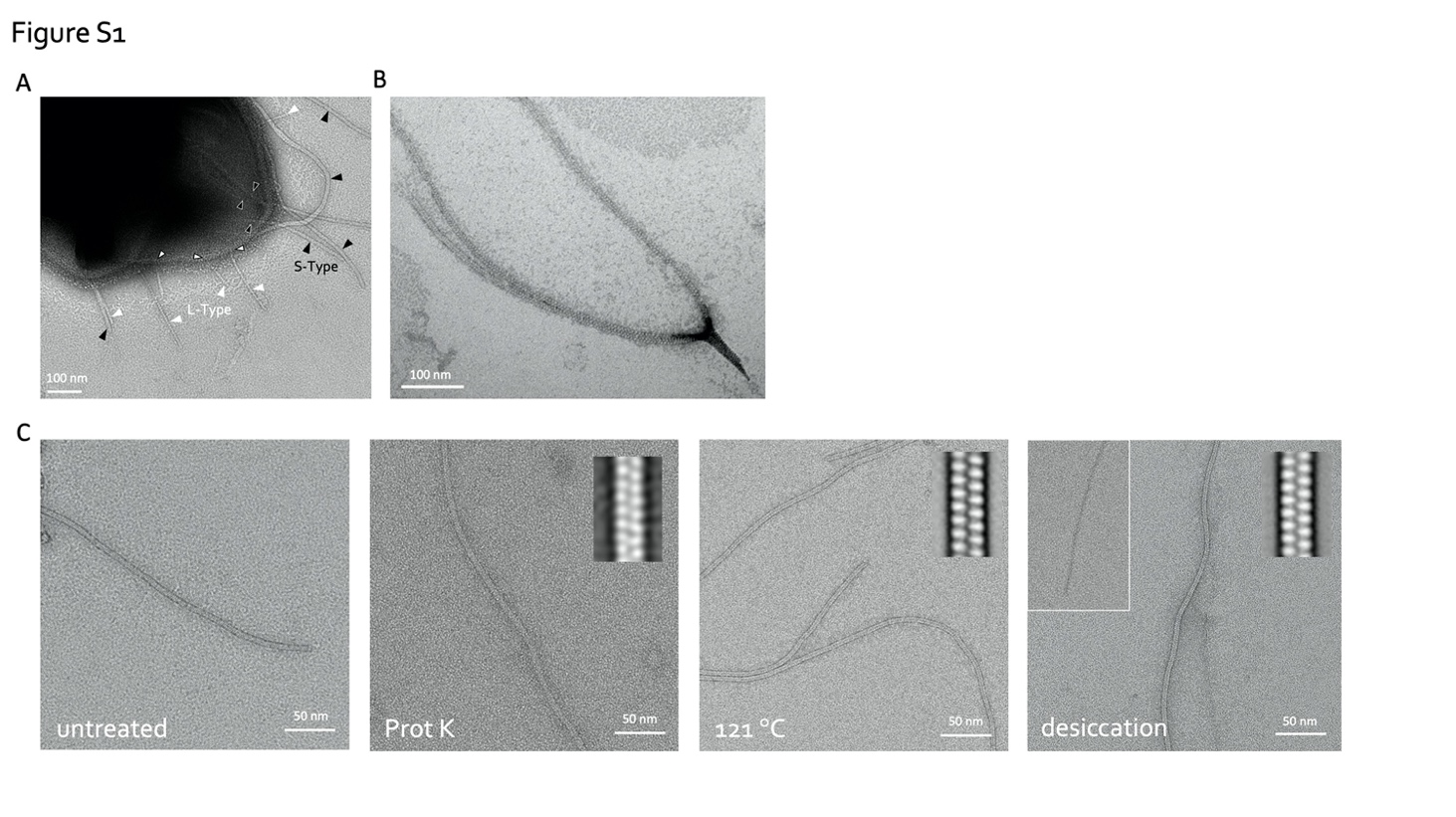

**Figure S1. Ena morphology and robustness. (A, B)** Negative stain TEM of *B. cereus* NVH 0075-95 endospore with indication of the two Ena morphologies: S-type (black arrowheads) and L-type Enas (white arrowheads) (A), and closed-up view of a dislodged S-type Ena bundle splitting into individual Ena fibers (B). **(C)** Negative stain TEM images of isolated *ex vivo* S-type Ena. To test Ena stability under different stresses, samples were treated, from left to right, with: (1) untreated control, (2) 1 hour of 1 mg/mL proteinase K, (3) autoclaving (i.e. 20 minutes at 121 °C) or (4) a 4 hour desiccation at 43 °C. Inset shows 2D class averages to assess the structural integrity of the treated Ena. S-type Enas are found to be resistant to Proteinase K treatment, autoclaving and desiccation at 43 °C, although some fibers appear to loose subunit integrity upon desiccation (inset). Desiccation at 43 °C may mimic conditions encountered by *Bacillus* spores during drought.

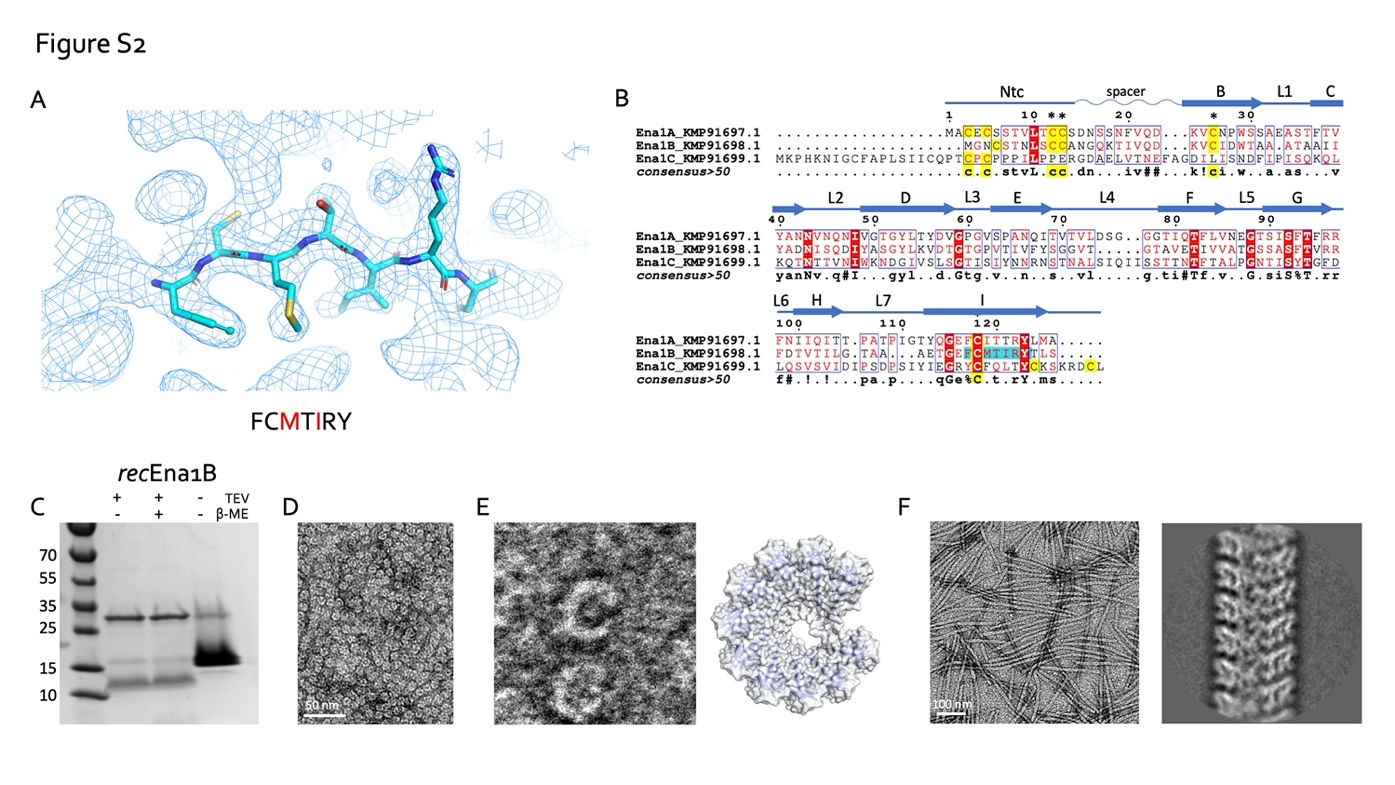

**Figure S2. S-type Ena structure determination and recombinant production.** (A) representative area of the 3D cryoEM potential map for *ex vivo* S-type Ena, at 3.2 Å resolution. An octameric peptide with sequence FCMTIRY was deduced *de novo* from the cryoEM potential map (shown in sticks) and used for a BLAST search of the *B. cereus* NVH 0075-95 genome. (B) Multiple sequence alignment of 3 ORF’s (KMP91697.1, KMP91698.1 and KMP91699.1) corresponding to DUF3992 containing proteins, of which the former two contain a sequence motif corresponding or similar to the one deduced from the EM potential map (shaded in cyan). The three ORFs are here shown to correspond to the S-type Ena subunits (see main text) and are hereafter referred to as Ena1A, Ena1B and Ena1C, respectively. Secondary structure and structural elements as determined from the built model (see Figure 2) are shown schematically above the sequences (Ntc: N-terminal connecter; arrows correspond to β-strands, labelled as in Figure 2). **(C)** SDS-PAGE of recombinant Ena1B, expressed in *E. coli*, affinity purified under denaturing conditions (8M urea) and treated with β-mercaptoethanol or TEV protease (to remove N-terminal 6xHis tag) as indicated. SDS-PAGE shows a species of apparent MW of ~12 and ~14 kDa, corresponding to the expected MW of the *rec*Ena1B monomer with and without the 6xHis tag removed by TEV digestion, respectively. **(D)** Negative stain TEM images of *rec*1Ena1B oligomers formed after refolding. **(E)** Closed up view that shows *rec*Ena1B oligomers form open crescents similar in dimensions and shape to single helical turns or arcs found in the S-type Ena fiber (model – right). Steric hindrance by the N-terminal His-tag is thought to arrest *rec*Ena1B polymerization into single helical arcs. **(F)** Negative stain image and 2D classification of Ena-like fibers formed after TEV digestion of *rec*Ena1B. Upon removal of the N-terminal His-tag, *rec*Ena1B readily assembles into fibers with helical properties closely resembling those found for *ex vivo* S-type Enas.

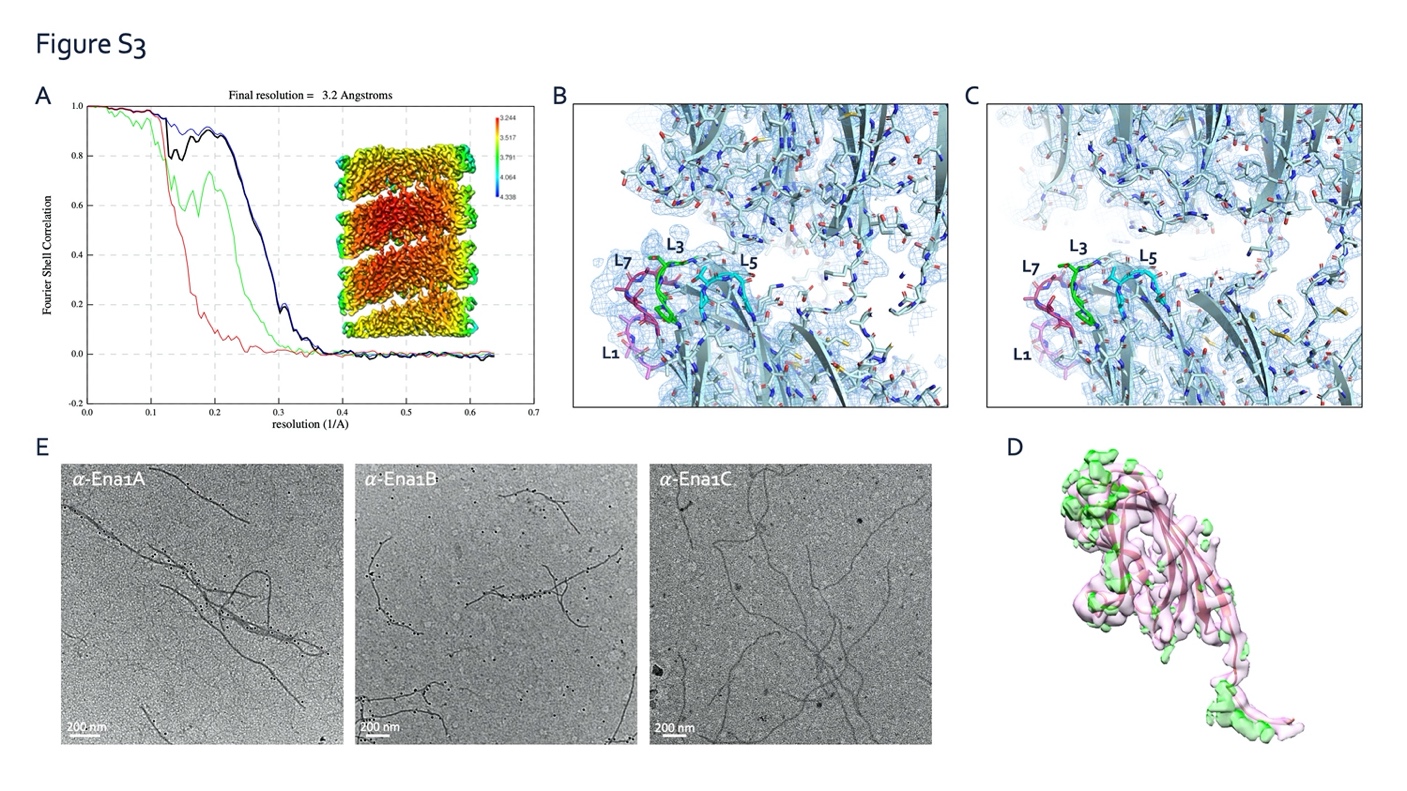

**Figure S3.** **S-type Ena are composed of both Ena1A and Ena1B subunits. (A)** FSC curve and local resolution heatmap (inset) of the *rec*Ena1B helical reconstruction, indicating a  final resolution of 3.2 Å at a cutoff of 0.143. FSC curve and local resolution were calculated by postprocessing in RELION3.0 using a solvent mask consisting of 3 helical turns. **(B, C)** Side-by-side comparison of cryo-EM maps calculated from of *ex vivo* (B) and *rec*ENA1B filaments (C), with the refined Ena1B model docked into the maps. The *ex vivo* Ena map shows features unaccounted for by the Ena1B model near loops 3 (L3) and 7 (L7), corresponding to regions of amino acid insertions in the Ena1A sequence (Figure S2B). **(D)** *rec*Ena1B map (pink) and *rec*Ena1B - *ex vivo* Ena1 difference map (green) masked over a single Ena1B subunit and calculated by TEMPy:Diffmap (Farabella et al., 2015) from the CCPEM package (Burnley et al., 2017). Difference in both maps locate to L3, L7 and the conformation of Ntc. **(E)** Immunogold TEM of *ex vivo* S-type Enas, stained with, from left to right, anti-Ena1A, anti-Ena1B and anti-Ena1C sera, each with gold-labeled (10 nm colloidal gold) anti-rabbit IgG as secondary antibody. Specific staining with Ena1A and Ena1B sera confirms the presence of both subunits in native Enas. No staining was seen with Ena1C serum.

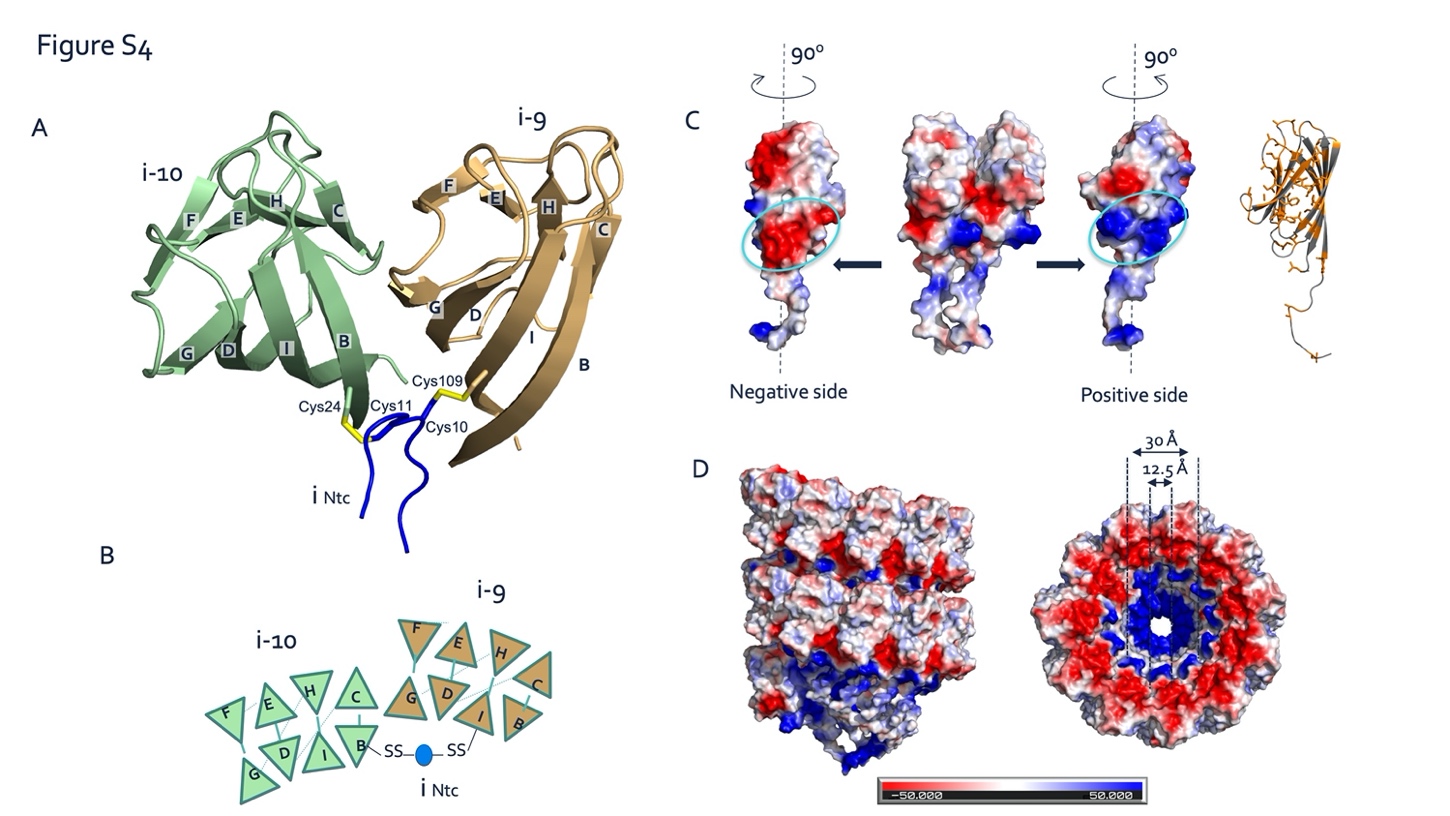

**Figure S4.** **Inter-subunit interactions in S-type Ena. (A, B)** Ribbon (A) and schematic (B) representation of lateral subunit – subunit contacts in S-type Ena. Strand G of BIDG sheet of each subunit is augmented with strand C of CHEF β-sheet of the succeeding subunit. Both subunits are covalently cross-linked via the Ntc (blue) of a subunit located, respectively, 9 or 10 subunits above. Cys11 and Cys10 go into a disulfide bond with residues 24 in the B strand of subunit i-10 and Cys109 in strand I of subunit i-9.  (**C, D**) Coulomb potential maps (calculated in PyMOL) of two adjacent subunits (C) and two helical turns of the S-type Ena showing the distribution of charge on the atomic model surface. Each subunit possesses complementary positive and negatively charged patches of residues at the inter-subunit surface that are responsible for electrostatic stabilizing interactions between the subunits. Similarly, stacked helical rings in the S-type Ena show a charge complementary interface (D).

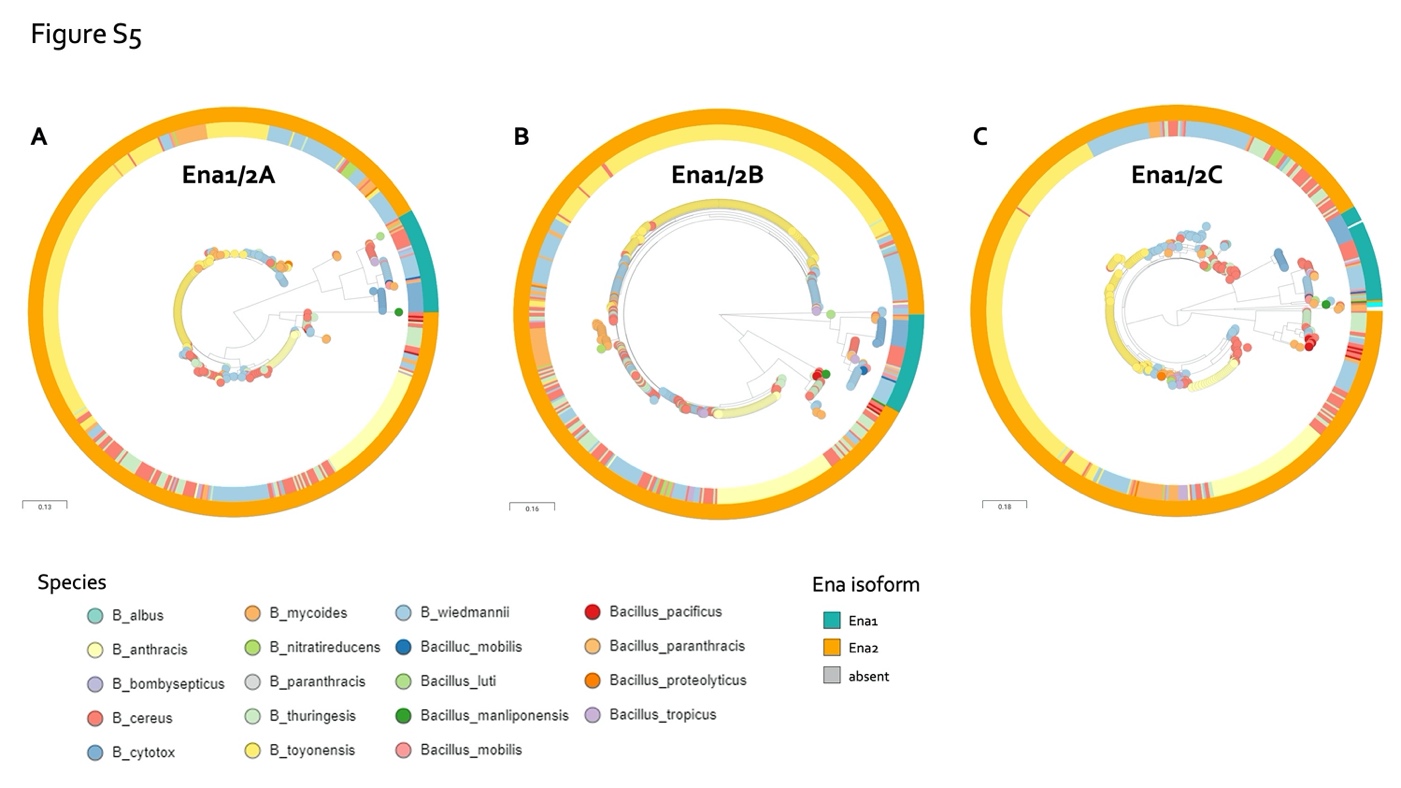

**Figure S5. Phylogenetic relationship between EnaA-C protein sequences among *Bacillus* spp.** Approximate likelihood trees generated by FastTree v.2.1.8 (Price et al., 2010), visualized in Microreact (Argimon et al., 2016). Trees are rooted on midpoint. Nodes are colored according to annotated species. See Methods for further details. **(A)** Relationship between Ena1A and Ena2A isoforms of 593 isolates. Ena1A and Ena2A are defined as ortho- or homologues having >90% coverage and >80% and 50-65% sequence identity, respectively, with Ena1A_GCF_001044825; KMP91697.1 protein sequence defined in Table S5. Interactive tree accessible at

<https://microreact.org/project/5UixxEY9vr2AVzXDVwa5t/1a8558fd>. **B)** Relationship between Ena1B and Ena2B isoforms of 591 isolates. Ena1B, Ena1B_candidate and Ena2B are defined as ortho- or homologues with >90% coverage and >80%, 60-80% and 40-60% sequence identity to Ena1B_NM_Oslo protein sequence defined in Table S5, respectively. Interactive tree accessible at <https://microreact.org/project/jJ4pARvqf9gyT916sTar5u/1332f3b3>. **(C)** Relationship between Ena1C and Ena2C isoforms of 591 isolates. Ena1C, Ena1C_candidate and Ena2C_candidates are defined as ortho- or homologues with >90% coverage and >80%, 60-80% and 40-60% sequence identity to Ena1C_NM_Oslo protein sequence defined in Table S5, respectively. Furthermore, isolates in which an ortho- or homologue was found elsewhere in the genome than the usual EnaA-B locus are colored cyan. Isolates that lacked an Ena1C homo- or orthologue are colored grey. Interactive tree accessible at <https://microreact.org/project/aQaqCUCJoj2mw55KQujbGY/099d7885>.

**Table S1** Cryo-EM model and data statistics

|  | *Ex vivo* S-type Ena  (EMD- 11592) | *rec*ENA1B  (EMD-11591)  (PDB 7A02) |
| --- | --- | --- |
| **Data collection and processing** | CryoARM300, BECM | CryoARM300, BECM |
| Magnification | 60.000 | 60.000 |
| Voltage (kV) | 300 | 300 |
| Electron exposure (e–/Å^2^) | 62.5 | 64.66 |
| Defocus range (μm) | -0.5 to -3.5 | -0.5 to -3.5 |
| Pixel size (Å) | 0.82 | 0.784 |
| Symmetry imposed | Helical  Rise= 3.22937  Rotation=31.0338 | Helical  Rise= 3.43721  Rotation=32.3504 |
| Initial particle images (no.) | 53501 | 100495 |
| Final particle images (no.) | 42822 | 65466 |
| Map resolution (Å)      FSC threshold | 3.2  0.143 | 3.05 0.143 |
| Map resolution range (Å) |  | 3.05-3.65 ^1^ |
| **Refinement** |  |  |
| Initial model used | NA | de novo |
| Model resolution (Å)      FSC threshold | NA  NA | 2.81  0.143 |
| Model resolution range (Å) |  |  |
| Map sharpening *B* factor (Å^2^) | 25.9 B-iso of density modification | 27.4 B-iso of density modification |
| Model composition      Non-hydrogen atoms      Protein residues      Ligands | NA  NA  NA | 18699 ^2^  2576 ^2^  0 |
| *B* factors (Å^2^)      Protein      Ligand | NA  NA | 54.39  NA |
| R.m.s. deviations      Bond lengths (Å)      Bond angles (°) | NA  NA | 0.008  0.736 |
| Validation      MolProbity score      Clashscore      Poor rotamers (%) | NA  NA  NA | 1.93  8.07  0 |
| Ramachandran plot      Favored (%)      Allowed (%)      Disallowed (%) | NA  NA  NA | 101 (92%) ^3^  9 (8%) ^3^  0 ^3^ |

^1^ Numbers reflect the density modified cryo-EM map calculated using ResolveCryoEM (Terwilliger et al., 2019)

^2^ Numbers reflect a S-type Ena model with 23 Ena1B protomers

^3^ Numbers for a single Ena1B protomer

**Table S2** Primers used in this work. To allow assembly of the PCR fragments, primers B and C contain sequences overlapping each other (italic).

| Primer | Sequence (5’-3’) |
| --- | --- |
| Deletion mutants | |
| *Δena1A* |  |
| A: 2184 | AATGGCGCCAGTTCAATTAC |
| B: 2198 | *CCTCTCTACATAGCCTT*TCCCCTCTCTCTT |
| C: 2199 | *AAGGCTATGTAGAGAGG*GGAATTAGTAT |
| D: 2178 | CCTCCTATTCTCCCACCTGAAA |
| *Δena1B* |  |
| A: 2164 | TCCATGTGGTATGGCAAAAA |
| B: 2165 | *CCATATATTACA****T****ACTAATT*CCCCTCTC |
| C: 2166 | *AATTAGTATGTAATATATGG*TGATTTAAAGATT |
| D: 2167 | AACCTACTTGCCCCTGTCCT |
| *Δena1C* |  |
| A: 2200 | CGCATCTTGTTTAGGTGCAA |
| B: 2201 | *ATTTTTTTGTTATCCTTTTCA*TAAGACTGTTTAC |
| C: 2202 | *TGAAAAGGATAACAAAAAAAT*TATTGCTTTTG |
| D: 2176 | AGGTGGAGGGACAATCCAAAC |
| *Δena1AB* |  |
| A: 2164 | TCCATGTGGTATGGCAAAAA |
| B: 2186 | *CCATATATTACATAGCCTTTCC*CCTCTC |
| C: 2197 | *AAAGGCTATGTAATATATGG*TGATTTAAAGAT |
| D: 2167 | AACCTACTTGCCCCTGTCCT |
| RT-PCR | |
| 2116/2117 | AAGTGCGTCTAATCAACAAGGAAA/ GGGAAATCTCCCATGAACACA |
| 2176/2177 | AGGTGGAGGGACAATCCAAAC/ GGCGAAACGTAAATGAAATGC |
| 2174/2175 | CCACTGGAAGTAGCGCATCTT / GCCGCTGTTCCAAGAATTGT |
| 2178/1279 | CCTCCTATTCTCCCACCTGAAA / CTCCAGCGAACTCATTGGTAACT |
| 2180/2181 | GGGTGTACGAGGGTGATATGAATT/ TGTCGTTCCGCCAAGTGTT |
| Complementation | |
| 2220/2221 | GCGGATGTTGTTGGACAA/ACGTGCAAACACATGAATCG |
| *rec*Ena1B | |
| Coding sequence | ATGCACCACCACCACCATCATTCTAGCGGTGAAAACCTGTACTTTCAGGGTAACTGCAGCACCAA  TCTGTCATGCTGTGCCAATGGTCAGAAGACCATTGTCCAGGATAAAGTCTGCATCGACTGGACCG  CAGCCGCTACTGCAGCAATCATTTACGCTGATAATATCAGCCAAGACATCTACGCTTCAGGCTATC  TGAAAGTGGATACAGGTACGGGTCCCGTGACCATCGTCTTTTACTCTGGTGGAGTCACAGGCACC  GCTGTGGAGACCATTGTGGTCGCCACGGGTTCGTCGGCCAGCTTTACGGTGCGCCGTTTTGATAC  CGTCACTATTCTGGGCACCGCAGCAGCGGAGACTGGTGAGTTTTGTATGACCATCCGTTACACTTT  GAGCTAA |
| Protein sequence | MHHHHHHSSGENLYFQGNCSTNLSCCANGQKTIVQDKVCIDWTAAATAAIIYADNISQDIYASGYLKV  DTGTGPVTIVFYSGGVTGTAVETIVVATGSSASFTVRRFDTVTILGTAAAETGEFCMTIRYTLS |

**Table S3** Overview of species included in the pairwise tBLASTn searches for the individual query proteins.

| Species | Number of genomes |
| --- | --- |
| *B. albus* | 1 |
| *B. anthracis* | 63 |
| *B. bombysepticus* | 1 |
| *B. cereus* | 85 |
| *B. cytotoxicus* | 14 |
| *B. manliponensis* | 1 |
| *B. gaemokensis* | 2 |
| *B. luti* | 1 |
| *B. mycoides* | 33 |
| *B. mobilis* | 5 |
| *B. nitratireducens* | 6 |
| *B. pacificus* | 3 |
| *B. paramycoides* | 2 |
| *B. paranthracis* | 3 |
| *B. pseudomycoides* | 8 |
| *B. proteolyticus* | 1 |
| *B. subtilis* | 127 |
| *B. thuringesis* | 50 |
| *B. toyonensis* | 204 |
| *B. tropicus* | 7 |
| *B. wiedmannii* | 119 |
| ***Total*** | **735** |

**Table S4** All genomes downloaded from RefSeq Database (*n=735*) and their annotated species, in addition to scaffolds/contigs of interest (112 *B. wiedermannii*, six *B. nitratireducens*, one *B. proteolyticus*, two *B. paramycoides*, five *B. pacificus*, five *B. mobilis*, 24 *B. mycoides*, one *B. manliponensis*, two *B. gaemokensis*). All strains were used in the pairwise tBLASTn search and to construct the k-mer based Mashtree.

| **Accession number** | **Species** |
| --- | --- |
| CP009335.1_genome | *B. thuringesis* |
| CP009720.1_genome | *B. thuringesis* |
| GCA_000171035.2_ASM17103v2_genomic.tsv | *B. cereus* |
| GCA_002952815.1_ASM295281v1_genomic.tsv | *B. cereus* |
| GCF_000003645.1_ASM364v1 | *B. cereus* |
| GCF_000003955.1_ASM395v1 | *B. cereus* |
| GCF_000007825.1_BcereusRef | *B. cereus* |
| GCF_000007845.1_ASM784v1 | *B. anthracis* |
| GCF_000008005.1_ASM800v1 | *B. cereus* |
| GCF_000008165.1_ASM816v1 | *B. anthracis* |
| GCF_000008445.1_ASM844v1 | *B. anthracis* |
| GCF_000008505.1_ASM850v1 | *B. thuringesis* |
| GCF_000009045.1_ASM904v1 | *B. subtilis* |
| GCF_000011625.1_ASM1162v1 | *B. cereus* |
| GCF_000013065.1_ASM1306v1 | *B. cereus* |
| GCF_000015065.1_ASM1506v1 | *B. thuringesis* |
| GCF_000017425.1_ASM1742v1 | *B. cytotox* |
| GCF_000021205.1_ASM2120v1 | *B. cereus* |
| GCF_000021225.1_ASM2122v1 | *B. cereus* |
| GCF_000021305.1_ASM2130v1 | *B. cereus* |
| GCF_000021445.1_ASM2144v1 | *B. anthracis* |
| GCF_000021785.1_ASM2178v1 | *B. cereus* |
| GCF_000022505.1_ASM2250v1 | *B. cereus* |
| GCF_000022865.1_ASM2286v1 | *B. anthracis* |
| GCF_000092165.1_ASM9216v1 | *B. thuringesis* |
| GCF_000143605.1_ASM14360v1 | *B. cereus* |
| GCF_000146565.1_ASM14656v1 | *B. subtilis* |
| GCF_000155325.1_ASM15532v1 | *B. subtilis* |
| GCF_000155355.1_ASM15535v1 | *B. subtilis* |
| GCF_000155375.1_ASM15537v1 | *B. subtilis* |
| GCF_000160895.1_ASM16089v1 | *B. cereus* |
| GCF_000160915.1_ASM16091v1 | *B. cereus* |
| GCF_000160935.1_ASM16093v1 | *B. cereus* |
| GCF_000160955.1_ASM16095v1 | *B. wiedmannii* |
| GCF_000160975.1_ASM16097v1 | *B. mycoides* |
| GCF_000161015.1_ASM16101v1 | *B. cereus* |
| GCF_000161035.1_ASM16103v1 | *B. cereus* |
| GCF_000161055.1_ASM16105v1 | *B. cereus* |
| GCF_000161075.1_ASM16107v1 | *B. cereus* |
| GCF_000161095.1_ASM16109v1 | *B. cereus* |
| GCF_000161115.1_ASM16111v1 | *B. cereus* |
| GCF_000161135.1_ASM16113v1 | *B. cereus* |
| GCF_000161155.1_ASM16115v1 | *B. cereus* |
| GCF_000161175.1_ASM16117v1 | *B. cereus* |
| GCF_000161195.1_ASM16119v1 | *B. cereus* |
| GCF_000161215.1_ASM16121v1 | *B. cereus* |
| GCF_000161235.1_ASM16123v1 | *B. cereus* |
| GCF_000161255.1_ASM16125v1 | *B. cereus* |
| GCF_000161275.1_ASM16127v1 | *B. cereus* |
| GCF_000161295.1_ASM16129v1 | *B. cereus* |
| GCF_000161315.1_ASM16131v1 | *B. cereus* |
| GCF_000161355.1_ASM16135v1 | *B. cereus* |
| GCF_000161375.1_ASM16137v1 | *B. cereus* |
| GCF_000161395.1_ASM16139v1 | *B. cereus* |
| GCF_000161415.1_ASM16141v1 | *B. pseudomycoides* |
| GCF_000161435.1_ASM16143v1 | *B. pseudomycoides* |
| GCF_000161455.1_ASM16145v1 | *B. pseudomycoides* |
| GCF_000181615.1_ASM18161v1 | *B. cereus* |
| GCF_000186085.1_ASM18608v1 | *B. subtilis* |
| GCF_000186745.1_ASM18674v1 | *B. subtilis* |
| GCF_000190515.1_ASM19051v1 | *B. thuringesis* |
| GCF_000193355.1_ASM19335v1 | *B. thuringesis* |
| GCF_000209795.2_ASM20979v2 | *B. subtilis* |
| GCF_000227465.1_ASM22746v1 | *B. subtilis* |
| GCF_000227485.1_ASM22748v1 | *B. subtilis* |
| GCF_000239195.1_ASM23919v1 | *B. cereus* |
| GCF_000258885.1_ASM25888v1 | *B. anthracis* |
| GCF_000283675.1_ASM28367v1 | *B. cereus* |
| GCF_000290695.1_Baci_cere_VDM022_V1 | *B. mycoides* |
| GCF_000290895.1_Baci_cere_VD078_V1 | *B. mycoides* |
| GCF_000290995.1_Baci_cere_AND1407_G13175_V1_genomic.tsv | *B. cereus* |
| GCF_000291015.1_Baci_cere_BAG1O-2_G13159_V1 | *B. toyonensis* |
| GCF_000291095.1_Baci_cere_HuB4-10_V1 | *B. toyonensis* |
| GCF_000291315.1_Baci_cere_CER074_V1 | *B. mycoides* |
| GCF_000291335.1_Baci_cere_CER057_V1 | *B. mycoides* |
| GCF_000291375.1_Baci_cere_BAG5X2-1_G13204_V1 | *B. wiedmannii* |
| GCF_000291495.1_Baci_cere_BAG2X1-2_G13195_V1 | *B. wiedmannii* |
| GCF_000292415.1_ASM29241v1 | *B. cereus* |
| GCF_000292455.1_ASM29245v1 | *B. thuringesis* |
| GCF_000292705.1_ASM29270v1 | *B. thuringesis* |
| GCF_000293565.1_Baci_cere_BAG6X1-1_G13210_V1 | *B. wiedmannii* |
| GCF_000293605.1_Baci_cere_BtB2-4_V1 | *B. mycoides* |
| GCF_000293765.1_ASM29376v1 | *B. subtilis* |
| GCF_000300475.1_ASM30047v1 | *B. thuringesis* |
| GCF_000306745.1_ASM30674v1 | *B. thuringesis* |
| GCF_000321395.1_ASM32139v1 | *B. subtilis* |
| GCF_000338315.1_BacF5 | *B. cereus* |
| GCF_000338735.1_ASM33873v1 | *B. subtilis* |
| GCF_000338755.1_ASM33875v1 | *B. thuringesis* |
| GCF_000341665.1_ASM34166v1 | *B. thuringesis* |
| GCF_000344745.1_ASM34474v1 | *B. subtilis* |
| GCF_000349795.1_ASM34979v1 | *B. subtilis* |
| GCF_000399045.1_Baci_cere_VDM019_V1 | *B. mycoides* |
| GCF_000496285.1_ASM49628v1 | *B. toyonensis* |
| GCF_000497485.1_ASM49748v1 | *B. subtilis* |
| GCF_000497525.1_ASM49752v2 | *B. thuringesis* |
| GCF_000512775.2_ASM51277v2 | *B. anthracis* |
| GCF_000512835.2_ASM51283v2 | *B. anthracis* |
| GCF_000523045.1_ASM52304v1 | *B. subtilis* |
| GCF_000534935.2_ASM53493v2 | *B. anthracis* |
| GCF_000558965.1_Ba8903G_1.0 | *B. anthracis* |
| GCF_000558985.1_Ba9080G_1.0 | *B. anthracis* |
| GCF_000559005.1_Ba52G_1.0 | *B. anthracis* |
| GCF_000583105.1_ASM58310v1 | *B. anthracis* |
| GCF_000635895.2_ASM63589v2 | *B. cereus* |
| GCF_000688795.1_ASM68879v1 | *B. thuringesis* |
| GCF_000699465.1_ASM69946v1 | *B. subtilis* |
| GCF_000699525.1_ASM69952v1 | *B. subtilis* |
| GCF_000706705.1_ASM70670v1 | *B. subtilis* |
| GCF_000712595.1_ASM71259v1 | *B. manliponensis* |
| GCF_000712615.1_ASM71261v1 | *B. gaemokensis* |
| GCF_000717535.1_ASM71753v1 | *B. thuringesis* |
| GCF_000725325.1_ASM72532v1 | *B. anthracis* |
| GCF_000737405.1_ASM73740v1 | *B. subtilis* |
| GCF_000742315.1_ASM74231v1 | *B. anthracis* |
| GCF_000742655.1_ASM74265v1 | *B. anthracis* |
| GCF_000742695.1_ASM74269v1 | *B. anthracis* |
| GCF_000742855.1_ASM74285v1 | *B. pseudomycoides* |
| GCF_000742875.1_ASM74287v1 | *B. anthracis* |
| GCF_000742895.1_ASM74289v1 | *B. anthracis* |
| GCF_000746965.1_BHU_1 | *B. pseudomycoides* |
| GCF_000747335.1_ASM74733v1 | *B. anthracis* |
| GCF_000747375.1_ASM74737v1 | *B. anthracis* |
| GCF_000747545.1_ASM74754v1 | *B. thuringesis* |
| GCF_000769515.1_ASM76951v1 | *B. subtilis* |
| GCF_000772125.1_ASM77212v1 | *B. subtilis* |
| GCF_000772165.1_ASM77216v1 | *B. subtilis* |
| GCF_000772205.1_ASM77220v1 | *B. subtilis* |
| GCF_000775975.1_ASM77597v1 | *B. mycoides* |
| GCF_000782835.1_ASM78283v1 | *B. subtilis* |
| GCF_000789275.1_ASM78927v1 | *B. subtilis* |
| GCF_000789295.1_ASM78929v1 | *B. subtilis* |
| GCF_000789315.1_ASM78931v1 | *B. cereus* |
| GCF_000803665.1_ASM80366v1 | *B. thuringesis* |
| GCF_000827065.1_ASM82706v1 | *B. subtilis* |
| GCF_000831065.1_ASM83106v1 | *B. bombysepticus* |
| GCF_000831505.1_ASM83150v1 | *B. anthracis* |
| GCF_000832385.1_ASM83238v1 | *B. cereus* |
| GCF_000832405.1_ASM83240v1 | *B. cereus* |
| GCF_000832425.1_ASM83242v1 | *B. anthracis* |
| GCF_000832445.1_ASM83244v1 | *B. anthracis* |
| GCF_000832465.1_ASM83246v1 | *B. anthracis* |
| GCF_000832485.1_ASM83248v1 | *B. thuringesis* |
| GCF_000832505.1_ASM83250v1 | *B. anthracis* |
| GCF_000832525.1_ASM83252v1 | *B. cereus* |
| GCF_000832565.1_ASM83256v1 | *B. anthracis* |
| GCF_000832585.1_ASM83258v1 | *B. anthracis* |
| GCF_000832605.1_ASM83260v1 | *B. mycoides* |
| GCF_000832635.1_ASM83263v1 | *B. anthracis* |
| GCF_000832665.1_ASM83266v1 | *B. anthracis* |
| GCF_000832725.1_ASM83272v1 | *B. anthracis* |
| GCF_000832745.1_ASM83274v1 | *B. anthracis* |
| GCF_000832765.1_ASM83276v1 | *B. cereus* |
| GCF_000832785.1_ASM83278v1 | *B. anthracis* |
| GCF_000832805.1_ASM83280v1 | *B. cereus* |
| GCF_000832825.1_ASM83282v1 | *B. thuringesis* |
| GCF_000832845.1_ASM83284v1 | *B. cereus* |
| GCF_000832865.1_ASM83286v1 | *B. cereus* |
| GCF_000832885.1_ASM83288v1 | *B. pseudomycoides* |
| GCF_000832925.1_ASM83292v1 | *B. thuringesis* |
| GCF_000832965.1_ASM83296v1 | *B. anthracis* |
| GCF_000833045.1_ASM83304v1 | *B. cereus* |
| GCF_000833065.1_ASM83306v1 | *B. anthracis* |
| GCF_000833085.1_ASM83308v1 | *B. thuringesis* |
| GCF_000833125.1_ASM83312v1 | *B. anthracis* |
| GCF_000833275.1_ASM83327v1 | *B. anthracis* |
| GCF_000835025.1_ASM83502v1 | *B. thuringesis* |
| GCF_000835185.1_ASM83518v1 | *B. cereus* |
| GCF_000875715.1_ASM87571v1 | *B. anthracis* |
| GCF_000931375.1_ASM93137v1 | *B. mycoides* |
| GCF_000940785.1_ASM94078v1 | *B. thuringesis* |
| GCF_000953615.1_BS49Ch | *B. subtilis* |
| GCF_000971925.1_ASM97192v1 | *B. subtilis* |
| GCF_000973605.1_ASM97360v1 | *B. subtilis* |
| GCF_000978375.1_ASM97837v1 | *B. cereus* |
| GCF_001008655.1_ASM100865v1 | *B. wiedmannii* |
| GCF_001015095.1_ASM101509v1 | *B. subtilis* |
| GCF_001017635.1_ASM101763v1 | *B. thuringesis* |
| GCF_001037985.1_ASM103798v1 | *B. subtilis* |
| GCF_001044575.1_ASM104457v1 | *B. wiedmannii* |
| GCF_001044745.1_ASM104474v1 | *B. wiedmannii* |
| GCF_001044775.1_ASM104477v1 | *B. toyonensis* |
| GCF_001044825.1_ASM104482v1 | *B. cereus* |
| GCF_001044935.1_ASM104493v1 | *B. mycoides* |
| GCF_001050335.1_ASM105033v1 | *B. cytotox* |
| GCF_001182785.1_ASM118278v1 | *B. thuringesis* |
| GCF_001183785.1_ASM118378v1 | *B. thuringesis* |
| GCF_001275045.2_ASM127504v2 | *B. toyonensis* |
| GCF_001277915.1_ASM127791v1 | *B. cereus* |
| GCF_001277955.1_ASM127795v1 | *B. anthracis* |
| GCF_001317525.1_CBN_33 | *B. wiedmannii* |
| GCF_001420855.1_ASM142085v1 | *B. thuringesis* |
| GCF_001455345.1_ASM145534v1 | *B. thuringesis* |
| GCF_001465815.1_ASM146581v1 | *B. subtilis* |
| GCF_001484805.1_M2E_15 | *B. mycoides* |
| GCF_001518875.1_ASM151887v1 | *B. cereus* |
| GCF_001534785.1_ASM153478v1 | *B. subtilis* |
| GCF_001541905.1_ASM154190v1 | *B. subtilis* |
| GCF_001543225.1_ASM154322v1 | *B. anthracis* |
| GCF_001548175.1_ASM154817v1 | *B. thuringesis* |
| GCF_001565875.1_ASM156587v1 | *B. subtilis* |
| GCF_001583695.1_ASM158369v1 | *B. wiedmannii* |
| GCF_001584035.1_ASM158403v1 | *B. wiedmannii* |
| GCF_001590835.1_ASM159083v1 | *B. gaemokensis* |
| GCF_001595725.1_ASM159572v1 | *B. thuringesis* |
| GCF_001596535.1_ASM159653v1 | *B. subtilis* |
| GCF_001597265.1_ASM159726v1 | *B. subtilis* |
| GCF_001598095.1_ASM159809v1 | *B. thuringesis* |
| GCF_001604995.1_ASM160499v1 | *B. subtilis* |
| GCF_001618665.1_ASM161866v1 | *B. thuringesis* |
| GCF_001619505.1_ASM161950v1 | *B. mycoides* |
| GCF_001635915.1_ASM163591v1 | *B. cereus* |
| GCF_001635955.1_ASM163595v1 | *B. cereus* |
| GCF_001635995.1_ASM163599v1 | *B. cereus* |
| GCF_001640965.1_ASM164096v1 | *B. thuringesis* |
| GCF_001645395.1_ASM164539v1 | *B. wiedmannii* |
| GCF_001645405.1_ASM164540v1 | *B. wiedmannii* |
| GCF_001645445.1_ASM164544v1 | *B. wiedmannii* |
| GCF_001645455.1_ASM164545v1 | *B. wiedmannii* |
| GCF_001645465.1_ASM164546v1 | *B. wiedmannii* |
| GCF_001645505.1_ASM164550v1 | *B. wiedmannii* |
| GCF_001645525.1_ASM164552v1 | *B. wiedmannii* |
| GCF_001645535.1_ASM164553v1 | *B. wiedmannii* |
| GCF_001645555.1_ASM164555v1 | *B. wiedmannii* |
| GCF_001654475.1_ASM165447v1 | *B. anthracis* |
| GCF_001660525.1_ASM166052v1 | *B. subtilis* |
| GCF_001675515.1_ASM167551v1 | *B. thuringesis* |
| GCF_001683065.1_ASM168306v1 | *B. anthracis* |
| GCF_001683095.1_ASM168309v1 | *B. anthracis* |
| GCF_001683135.1_ASM168313v1 | *B. anthracis* |
| GCF_001683155.1_ASM168315v1 | *B. anthracis* |
| GCF_001683175.1_ASM168317v1 | *B. anthracis* |
| GCF_001683195.1_ASM168319v1 | *B. anthracis* |
| GCF_001683215.1_ASM168321v1 | *B. anthracis* |
| GCF_001683235.1_ASM168323v1 | *B. anthracis* |
| GCF_001683255.1_ASM168325v1 | *B. anthracis* |
| GCF_001683275.1_ASM168327v1 | *B. anthracis* |
| GCF_001683295.1_ASM168329v1 | *B. anthracis* |
| GCF_001685565.1_ASM168556v1 | *B. thuringesis* |
| GCF_001692675.1_ASM169267v1 | *B. thuringesis* |
| GCF_001697265.1_ASM169726v1 | *B. subtilis* |
| GCF_001703495.1_ASM170349v1 | *B. subtilis* |
| GCF_001704095.1_ASM170409v1 | *B. subtilis* |
| GCF_001720505.1_ASM172050v1 | *B. subtilis* |
| GCF_001721145.1_ASM172114v1 | *B. cereus* |
| GCF_001721165.1_ASM172116v1 | *B. thuringesis* |
| GCF_001746575.1_ASM174657v1 | *B. subtilis* |
| GCF_001747445.1_ASM174744v1 | *B. subtilis* |
| GCF_001750745.1_ASM175074v1 | *B. subtilis* |
| GCF_001756265.1_ASM175626v1 | *B. wiedmannii* |
| GCF_001808235.1_ASM180823v1 | *B. subtilis* |
| GCF_001880305.1_ASM188030v1 | *B. cereus* |
| GCF_001884025.1_ASM188402v1 | *B. pacificus* |
| GCF_001884035.1_ASM188403v1 | *B. tropicus* |
| GCF_001884045.1_ASM188404v1 | *B. mobilis* |
| GCF_001884065.1_ASM188406v1 | *B. proteolyticus* |
| GCF_001884105.1_ASM188410v1 | *B. luti* |
| GCF_001884135.1_ASM188413v1 | *B. nitratireducens* |
| GCF_001884235.1_ASM188423v1 | *B. paramycoides* |
| GCF_001889385.1_ASM188938v1 | *B. subtilis* |
| GCF_001889625.1_ASM188962v1 | *B. subtilis* |
| GCF_001890405.1_ASM189040v1 | *B. subtilis* |
| GCF_001902555.1_ASM190255v1 | *B. subtilis* |
| GCF_001932005.1_ASM193200v1 | *B. wiedmannii* |
| GCF_001936375.1_ASM193637v1 | *B. anthracis* |
| GCF_001941885.1_ASM194188v1 | *B. cereus* |
| GCF_001941905.1_ASM194190v1 | *B. cereus* |
| GCF_001941925.1_ASM194192v1 | *B. cereus* |
| GCF_001990245.1_ASM199024v1 | *B. anthracis* |
| GCF_002000005.1_ASM200000v1 | *B. cereus* |
| GCF_002005265.1_ASM200526v1 | *B. anthracis* |
| GCF_002009095.1_ASM200909v1 | *B. subtilis* |
| GCF_002009135.1_ASM200913v1 | *B. subtilis* |
| GCF_002014615.1_ASM201461v1 | *B. pseudomycoides* |
| GCF_002014635.1_ASM201463v1 | *B. wiedmannii* |
| GCF_002025105.1_ASM202510v1 | *B. thuringesis* |
| GCF_002055965.1_ASM205596v1 | *B. subtilis* |
| GCF_002071845.1_ASM207184v1 | *B. toyonensis* |
| GCF_002072735.1_ASM207273v1 | *B. subtilis* |
| GCF_002096095.1_ASM209609v1 | *B. subtilis* |
| GCF_002104575.1_ASM210457v1 | *B. mycoides* |
| GCF_002109455.1_ASM210945v1 | *B. toyonensis* |
| GCF_002117465.1_ASM211746v1 | *B. cereus* |
| GCF_002118205.1_ASM211820v1 | *B. mycoides* |
| GCF_002119445.1_ASM211944v1 | *B. thuringesis* |
| GCF_002142595.1_ASM214259v1 | *B. subtilis* |
| GCF_002147065.1_ASM214706v1 | *B. wiedmannii* |
| GCF_002158195.1_ASM215819v1 | *B. pseudomycoides* |
| GCF_002163815.1_ASM216381v1 | *B. subtilis* |
| GCF_002173615.1_ASM217361v1 | *B. subtilis* |
| GCF_002173695.1_ASM217369v1 | *B. subtilis* |
| GCF_002173715.1_ASM217371v1 | *B. subtilis* |
| GCF_002173755.1_ASM217375v1 | *B. thuringesis* |
| GCF_002184245.1_ASM218424v1 | *B. thuringesis* |
| GCF_002192515.1_ASM219251v1 | *B. thuringesis* |
| GCF_002200005.1_ASM220000v1 | *B. mycoides* |
| GCF_002200135.1_ASM220013v1 | *B. mycoides* |
| GCF_002200215.1_ASM220021v1 | *B. wiedmannii* |
| GCF_002201955.1_ASM220195v1 | *B. subtilis* |
| GCF_002201995.1_ASM220199v1 | *B. subtilis* |
| GCF_002202035.1_ASM220203v1 | *B. subtilis* |
| GCF_002202055.1_ASM220205v1 | *B. subtilis* |
| GCF_002205435.1_ASM220543v1 | *B. subtilis* |
| GCF_002208785.1_ASM220878v2 | *B. anthracis* |
| GCF_002214705.1_ASM221470v1 | *B. cereus* |
| GCF_002214725.1_ASM221472v1 | *B. cereus* |
| GCF_002214765.1_ASM221476v1 | *B. cereus* |
| GCF_002215175.1_ASM221517v1 | *B. cereus* |
| GCF_002216085.1_ASM221608v1 | *B. subtilis* |
| GCF_002216125.1_ASM221612v1 | *B. cereus* |
| GCF_002220075.1_ASM222007v1 | *B. subtilis* |
| GCF_002220285.1_ASM222028v1 | *B. cereus* |
| GCF_002222555.1_ASM222255v1 | *B. thuringesis* |
| GCF_002224345.1_ASM222434v1 | *B. cereus* |
| GCF_002250885.2_ASM225088v2 | *B. cytotox* |
| GCF_002250905.2_ASM225090v2 | *B. cytotox* |
| GCF_002250925.2_ASM225092v2 | *B. cytotox* |
| GCF_002250945.2_ASM225094v2 | *B. cytotox* |
| GCF_002250965.2_ASM225096v2 | *B. cytotox* |
| GCF_002251005.2_ASM225100v2 | *B. cytotox* |
| GCF_002251025.2_ASM225102v2 | *B. cytotox* |
| GCF_002251045.2_ASM225104v2 | *B. cytotox* |
| GCF_002251055.2_ASM225105v2 | *B. cytotox* |
| GCF_002251115.2_ASM225111v2 | *B. cytotox* |
| GCF_002269175.1_ASM226917v1 | *B. subtilis* |
| GCF_002269195.1_ASM226919v1 | *B. subtilis* |
| GCF_002277915.1_ASM227791v1 | *B. anthracis* |
| GCF_002290105.1_ASM229010v1 | *B. cereus* |
| GCF_002290305.1_ASM229030v1 | *B. subtilis* |
| GCF_002356575.1_ASM235657v1 | *B. anthracis* |
| GCF_002550225.1_ASM255022v1 | *B. wiedmannii* |
| GCF_002550285.1_ASM255028v1 | *B. toyonensis* |
| GCF_002550355.1_ASM255035v1 | *B. toyonensis* |
| GCF_002550525.1_ASM255052v1 | *B. toyonensis* |
| GCF_002550545.1_ASM255054v1 | *B. toyonensis* |
| GCF_002550635.1_ASM255063v1 | *B. wiedmannii* |
| GCF_002550725.1_ASM255072v1 | *B. toyonensis* |
| GCF_002550875.1_ASM255087v1 | *B. wiedmannii* |
| GCF_002550925.1_ASM255092v1 | *B. toyonensis* |
| GCF_002550975.1_ASM255097v1 | *B. toyonensis* |
| GCF_002551545.1_ASM255154v1 | *B. wiedmannii* |
| GCF_002551565.1_ASM255156v1 | *B. toyonensis* |
| GCF_002551705.1_ASM255170v1 | *B. toyonensis* |
| GCF_002551725.1_ASM255172v1 | *B. toyonensis* |
| GCF_002551735.1_ASM255173v1 | *B. toyonensis* |
| GCF_002551965.1_ASM255196v1 | *B. toyonensis* |
| GCF_002552055.1_ASM255205v1 | *B. toyonensis* |
| GCF_002552125.1_ASM255212v1 | *B. toyonensis* |
| GCF_002552205.1_ASM255220v1 | *B. wiedmannii* |
| GCF_002552225.1_ASM255222v1 | *B. toyonensis* |
| GCF_002552405.1_ASM255240v1 | *B. toyonensis* |
| GCF_002552545.1_ASM255254v1 | *B. toyonensis* |
| GCF_002552615.1_ASM255261v1 | *B. toyonensis* |
| GCF_002552675.1_ASM255267v1 | *B. toyonensis* |
| GCF_002552685.1_ASM255268v1 | *B. toyonensis* |
| GCF_002552775.1_ASM255277v1 | *B. toyonensis* |
| GCF_002552785.1_ASM255278v1 | *B. toyonensis* |
| GCF_002552975.1_ASM255297v1 | *B. toyonensis* |
| GCF_002553125.1_ASM255312v1 | *B. wiedmannii* |
| GCF_002553325.1_ASM255332v1 | *B. toyonensis* |
| GCF_002553375.1_ASM255337v1 | *B. toyonensis* |
| GCF_002553405.1_ASM255340v1 | *B. wiedmannii* |
| GCF_002553415.1_ASM255341v1 | *B. toyonensis* |
| GCF_002554755.1_ASM255475v1 | *B. toyonensis* |
| GCF_002554835.1_ASM255483v1 | *B. wiedmannii* |
| GCF_002554865.1_ASM255486v1 | *B. toyonensis* |
| GCF_002554905.1_ASM255490v1 | *B. toyonensis* |
| GCF_002554925.1_ASM255492v1 | *B. toyonensis* |
| GCF_002554935.1_ASM255493v1 | *B. mycoides* |
| GCF_002554965.1_ASM255496v1 | *B. toyonensis* |
| GCF_002554975.1_ASM255497v1 | *B. wiedmannii* |
| GCF_002555025.1_ASM255502v1 | *B. toyonensis* |
| GCF_002555035.1_ASM255503v1 | *B. toyonensis* |
| GCF_002555055.1_ASM255505v1 | *B. toyonensis* |
| GCF_002555085.1_ASM255508v1 | *B. wiedmannii* |
| GCF_002555095.1_ASM255509v1 | *B. wiedmannii* |
| GCF_002555115.1_ASM255511v1 | *B. toyonensis* |
| GCF_002555155.1_ASM255515v1 | *B. wiedmannii* |
| GCF_002555165.1_ASM255516v1 | *B. wiedmannii* |
| GCF_002555225.1_ASM255522v1 | *B. toyonensis* |
| GCF_002555265.1_ASM255526v1 | *B. wiedmannii* |
| GCF_002555305.1_ASM255530v1 | *B. wiedmannii* |
| GCF_002555315.1_ASM255531v1 | *B. toyonensis* |
| GCF_002555345.1_ASM255534v1 | *B. toyonensis* |
| GCF_002555385.1_ASM255538v1 | *B. wiedmannii* |
| GCF_002555405.1_ASM255540v1 | *B. toyonensis* |
| GCF_002555465.1_ASM255546v1 | *B. toyonensis* |
| GCF_002555485.1_ASM255548v1 | *B. wiedmannii* |
| GCF_002555495.1_ASM255549v1 | *B. toyonensis* |
| GCF_002555505.1_ASM255550v1 | *B. wiedmannii* |
| GCF_002555555.1_ASM255555v1 | *B. wiedmannii* |
| GCF_002555585.1_ASM255558v1 | *B. toyonensis* |
| GCF_002555605.1_ASM255560v1 | *B. toyonensis* |
| GCF_002555645.1_ASM255564v1 | *B. wiedmannii* |
| GCF_002555665.1_ASM255566v1 | *B. wiedmannii* |
| GCF_002555735.1_ASM255573v1 | *B. wiedmannii* |
| GCF_002555765.1_ASM255576v1 | *B. toyonensis* |
| GCF_002555805.1_ASM255580v1 | *B. toyonensis* |
| GCF_002555845.1_ASM255584v1 | *B. toyonensis* |
| GCF_002555855.1_ASM255585v1 | *B. wiedmannii* |
| GCF_002555885.1_ASM255588v1 | *B. wiedmannii* |
| GCF_002555905.1_ASM255590v1 | *B. toyonensis* |
| GCF_002555915.1_ASM255591v1 | *B. toyonensis* |
| GCF_002555945.1_ASM255594v1 | *B. wiedmannii* |
| GCF_002555955.1_ASM255595v1 | *B. toyonensis* |
| GCF_002555975.1_ASM255597v1 | *B. toyonensis* |
| GCF_002556015.1_ASM255601v1 | *B. toyonensis* |
| GCF_002556065.1_ASM255606v1 | *B. wiedmannii* |
| GCF_002556085.1_ASM255608v1 | *B. wiedmannii* |
| GCF_002556095.1_ASM255609v1 | *B. wiedmannii* |
| GCF_002556125.1_ASM255612v1 | *B. toyonensis* |
| GCF_002556135.1_ASM255613v1 | *B. toyonensis* |
| GCF_002556155.1_ASM255615v1 | *B. toyonensis* |
| GCF_002556175.1_ASM255617v1 | *B. wiedmannii* |
| GCF_002556225.1_ASM255622v1 | *B. toyonensis* |
| GCF_002556265.1_ASM255626v1 | *B. toyonensis* |
| GCF_002556275.1_ASM255627v1 | *B. wiedmannii* |
| GCF_002556325.1_ASM255632v1 | *B. toyonensis* |
| GCF_002556405.1_ASM255640v1 | *B. wiedmannii* |
| GCF_002556425.1_ASM255642v1 | *B. wiedmannii* |
| GCF_002556435.1_ASM255643v1 | *B. toyonensis* |
| GCF_002556675.1_ASM255667v1 | *B. toyonensis* |
| GCF_002556705.1_ASM255670v1 | *B. toyonensis* |
| GCF_002556715.1_ASM255671v1 | *B. toyonensis* |
| GCF_002556745.1_ASM255674v1 | *B. wiedmannii* |
| GCF_002556765.1_ASM255676v1 | *B. toyonensis* |
| GCF_002556775.1_ASM255677v1 | *B. toyonensis* |
| GCF_002556825.1_ASM255682v1 | *B. wiedmannii* |
| GCF_002556835.1_ASM255683v1 | *B. toyonensis* |
| GCF_002556885.1_ASM255688v1 | *B. toyonensis* |
| GCF_002556905.1_ASM255690v1 | *B. wiedmannii* |
| GCF_002556915.1_ASM255691v1 | *B. wiedmannii* |
| GCF_002556945.1_ASM255694v1 | *B. toyonensis* |
| GCF_002556955.1_ASM255695v1 | *B. toyonensis* |
| GCF_002557005.1_ASM255700v1 | *B. toyonensis* |
| GCF_002557015.1_ASM255701v1 | *B. wiedmannii* |
| GCF_002557045.1_ASM255704v1 | *B. toyonensis* |
| GCF_002557085.1_ASM255708v1 | *B. wiedmannii* |
| GCF_002557115.1_ASM255711v1 | *B. toyonensis* |
| GCF_002557145.1_ASM255714v1 | *B. toyonensis* |
| GCF_002557155.1_ASM255715v1 | *B. wiedmannii* |
| GCF_002557185.1_ASM255718v1 | *B. wiedmannii* |
| GCF_002557205.1_ASM255720v1 | *B. wiedmannii* |
| GCF_002557225.1_ASM255722v1 | *B. toyonensis* |
| GCF_002557255.1_ASM255725v1 | *B. toyonensis* |
| GCF_002557295.1_ASM255729v1 | *B. wiedmannii* |
| GCF_002557315.1_ASM255731v1 | *B. wiedmannii* |
| GCF_002557345.1_ASM255734v1 | *B. wiedmannii* |
| GCF_002557355.1_ASM255735v1 | *B. wiedmannii* |
| GCF_002557405.1_ASM255740v1 | *B. wiedmannii* |
| GCF_002557445.1_ASM255744v1 | *B. toyonensis* |
| GCF_002557475.1_ASM255747v1 | *B. wiedmannii* |
| GCF_002557505.1_ASM255750v1 | *B. toyonensis* |
| GCF_002557515.1_ASM255751v1 | *B. toyonensis* |
| GCF_002557555.1_ASM255755v1 | *B. toyonensis* |
| GCF_002557595.1_ASM255759v1 | *B. wiedmannii* |
| GCF_002557665.1_ASM255766v1 | *B. wiedmannii* |
| GCF_002557675.1_ASM255767v1 | *B. wiedmannii* |
| GCF_002557845.1_ASM255784v1 | *B. wiedmannii* |
| GCF_002557855.1_ASM255785v1 | *B. wiedmannii* |
| GCF_002557865.1_ASM255786v1 | *B. toyonensis* |
| GCF_002568685.1_ASM256868v1 | *B. wiedmannii* |
| GCF_002568725.1_ASM256872v1 | *B. toyonensis* |
| GCF_002568745.1_ASM256874v1 | *B. toyonensis* |
| GCF_002568785.1_ASM256878v1 | *B. wiedmannii* |
| GCF_002568805.1_ASM256880v1 | *B. wiedmannii* |
| GCF_002568825.1_ASM256882v1 | *B. toyonensis* |
| GCF_002568845.1_ASM256884v1 | *B. toyonensis* |
| GCF_002568855.1_ASM256885v1 | *B. toyonensis* |
| GCF_002568865.1_ASM256886v1 | *B. toyonensis* |
| GCF_002568875.1_ASM256887v1 | *B. wiedmannii* |
| GCF_002568925.1_ASM256892v1 | *B. wiedmannii* |
| GCF_002568935.1_ASM256893v1 | *B. mycoides* |
| GCF_002568955.1_ASM256895v1 | *B. wiedmannii* |
| GCF_002569035.1_ASM256903v1 | *B. toyonensis* |
| GCF_002569065.1_ASM256906v1 | *B. toyonensis* |
| GCF_002569095.1_ASM256909v1 | *B. toyonensis* |
| GCF_002569115.1_ASM256911v1 | *B. toyonensis* |
| GCF_002569125.1_ASM256912v1 | *B. wiedmannii* |
| GCF_002569165.1_ASM256916v1 | *B. wiedmannii* |
| GCF_002569175.1_ASM256917v1 | *B. toyonensis* |
| GCF_002569215.1_ASM256921v1 | *B. wiedmannii* |
| GCF_002569295.1_ASM256929v1 | *B. toyonensis* |
| GCF_002569325.1_ASM256932v1 | *B. toyonensis* |
| GCF_002569365.1_ASM256936v1 | *B. toyonensis* |
| GCF_002569385.1_ASM256938v1 | *B. mycoides* |
| GCF_002569395.1_ASM256939v1 | *B. toyonensis* |
| GCF_002569425.1_ASM256942v1 | *B. toyonensis* |
| GCF_002569445.1_ASM256944v1 | *B. toyonensis* |
| GCF_002569455.1_ASM256945v1 | *B. toyonensis* |
| GCF_002569465.1_ASM256946v1 | *B. toyonensis* |
| GCF_002569495.1_ASM256949v1 | *B. toyonensis* |
| GCF_002569515.1_ASM256951v1 | *B. toyonensis* |
| GCF_002569525.1_ASM256952v1 | *B. toyonensis* |
| GCF_002569595.1_ASM256959v1 | *B. wiedmannii* |
| GCF_002569665.1_ASM256966v1 | *B. wiedmannii* |
| GCF_002569675.1_ASM256967v1 | *B. wiedmannii* |
| GCF_002569715.1_ASM256971v1 | *B. wiedmannii* |
| GCF_002569745.1_ASM256974v1 | *B. toyonensis* |
| GCF_002569805.1_ASM256980v1 | *B. toyonensis* |
| GCF_002569835.1_ASM256983v1 | *B. toyonensis* |
| GCF_002569895.1_ASM256989v1 | *B. toyonensis* |
| GCF_002569955.1_ASM256995v1 | *B. toyonensis* |
| GCF_002569995.1_ASM256999v1 | *B. wiedmannii* |
| GCF_002570025.1_ASM257002v1 | *B. toyonensis* |
| GCF_002570035.1_ASM257003v1 | *B. wiedmannii* |
| GCF_002570045.1_ASM257004v1 | *B. toyonensis* |
| GCF_002570065.1_ASM257006v1 | *B. toyonensis* |
| GCF_002570105.1_ASM257010v1 | *B. toyonensis* |
| GCF_002570115.1_ASM257011v1 | *B. toyonensis* |
| GCF_002570135.1_ASM257013v1 | *B. wiedmannii* |
| GCF_002570155.1_ASM257015v1 | *B. toyonensis* |
| GCF_002570165.1_ASM257016v1 | *B. toyonensis* |
| GCF_002570205.1_ASM257020v1 | *B. toyonensis* |
| GCF_002570235.1_ASM257023v1 | *B. toyonensis* |
| GCF_002570255.1_ASM257025v1 | *B. wiedmannii* |
| GCF_002570265.1_ASM257026v1 | *B. wiedmannii* |
| GCF_002570305.1_ASM257030v1 | *B. wiedmannii* |
| GCF_002570335.1_ASM257033v1 | *B. wiedmannii* |
| GCF_002570345.1_ASM257034v1 | *B. toyonensis* |
| GCF_002570385.1_ASM257038v1 | *B. toyonensis* |
| GCF_002570405.1_ASM257040v1 | *B. toyonensis* |
| GCF_002571435.1_ASM257143v1 | *B. toyonensis* |
| GCF_002571465.1_ASM257146v1 | *B. wiedmannii* |
| GCF_002571525.1_ASM257152v1 | *B. wiedmannii* |
| GCF_002571555.1_ASM257155v1 | *B. toyonensis* |
| GCF_002571605.1_ASM257160v1 | *B. toyonensis* |
| GCF_002571615.1_ASM257161v1 | *B. toyonensis* |
| GCF_002571635.1_ASM257163v1 | *B. wiedmannii* |
| GCF_002571675.1_ASM257167v1 | *B. toyonensis* |
| GCF_002571685.1_ASM257168v1 | *B. wiedmannii* |
| GCF_002571695.1_ASM257169v1 | *B. toyonensis* |
| GCF_002571725.1_ASM257172v1 | *B. wiedmannii* |
| GCF_002571765.1_ASM257176v1 | *B. toyonensis* |
| GCF_002571785.1_ASM257178v1 | *B. wiedmannii* |
| GCF_002571795.1_ASM257179v1 | *B. wiedmannii* |
| GCF_002571825.1_ASM257182v1 | *B. toyonensis* |
| GCF_002571885.1_ASM257188v1 | *B. toyonensis* |
| GCF_002571915.1_ASM257191v1 | *B. toyonensis* |
| GCF_002571935.1_ASM257193v1 | *B. toyonensis* |
| GCF_002571965.1_ASM257196v1 | *B. toyonensis* |
| GCF_002572005.1_ASM257200v1 | *B. toyonensis* |
| GCF_002572035.1_ASM257203v1 | *B. toyonensis* |
| GCF_002572045.1_ASM257204v1 | *B. wiedmannii* |
| GCF_002572105.1_ASM257210v1 | *B. wiedmannii* |
| GCF_002572145.1_ASM257214v1 | *B. wiedmannii* |
| GCF_002572155.1_ASM257215v1 | *B. toyonensis* |
| GCF_002572165.1_ASM257216v1 | *B. toyonensis* |
| GCF_002572225.1_ASM257222v1 | *B. wiedmannii* |
| GCF_002572235.1_ASM257223v1 | *B. toyonensis* |
| GCF_002572245.1_ASM257224v1 | *B. toyonensis* |
| GCF_002572295.1_ASM257229v1 | *B. toyonensis* |
| GCF_002572315.1_ASM257231v1 | *B. wiedmannii* |
| GCF_002572325.1_ASM257232v1 | *B. wiedmannii* |
| GCF_002572395.1_ASM257239v1 | *B. toyonensis* |
| GCF_002572415.1_ASM257241v1 | *B. toyonensis* |
| GCF_002572455.1_ASM257245v1 | *B. toyonensis* |
| GCF_002572475.1_ASM257247v1 | *B. toyonensis* |
| GCF_002573545.1_ASM257354v1 | *B. toyonensis* |
| GCF_002578845.1_ASM257884v1 | *B. toyonensis* |
| GCF_002578895.1_ASM257889v1 | *B. wiedmannii* |
| GCF_002578965.1_ASM257896v1 | *B. toyonensis* |
| GCF_002578975.1_ASM257897v1 | *B. wiedmannii* |
| GCF_002579095.1_ASM257909v1 | *B. wiedmannii* |
| GCF_002579245.1_ASM257924v1 | *B. toyonensis* |
| GCF_002579305.1_ASM257930v1 | *B. toyonensis* |
| GCF_002579355.1_ASM257935v1 | *B. toyonensis* |
| GCF_002579445.1_ASM257944v1 | *B. wiedmannii* |
| GCF_002579545.1_ASM257954v1 | *B. wiedmannii* |
| GCF_002579615.1_ASM257961v1 | *B. toyonensis* |
| GCF_002579625.1_ASM257962v1 | *B. toyonensis* |
| GCF_002579725.1_ASM257972v1 | *B. wiedmannii* |
| GCF_002579735.1_ASM257973v1 | *B. toyonensis* |
| GCF_002579775.1_ASM257977v1 | *B. toyonensis* |
| GCF_002579865.1_ASM257986v1 | *B. wiedmannii* |
| GCF_002579895.1_ASM257989v1 | *B. toyonensis* |
| GCF_002579945.1_ASM257994v1 | *B. wiedmannii* |
| GCF_002580025.1_ASM258002v1 | *B. wiedmannii* |
| GCF_002580085.1_ASM258008v1 | *B. toyonensis* |
| GCF_002580095.1_ASM258009v1 | *B. wiedmannii* |
| GCF_002580125.1_ASM258012v1 | *B. toyonensis* |
| GCF_002580305.1_ASM258030v1 | *B. toyonensis* |
| GCF_002580355.1_ASM258035v1 | *B. toyonensis* |
| GCF_002580365.1_ASM258036v1 | *B. toyonensis* |
| GCF_002580455.1_ASM258045v1 | *B. toyonensis* |
| GCF_002580465.1_ASM258046v1 | *B. toyonensis* |
| GCF_002580575.1_ASM258057v1 | *B. toyonensis* |
| GCF_002580725.1_ASM258072v1 | *B. toyonensis* |
| GCF_002580865.1_ASM258086v1 | *B. toyonensis* |
| GCF_002580895.1_ASM258089v1 | *B. toyonensis* |
| GCF_002580965.1_ASM258096v1 | *B. toyonensis* |
| GCF_002580975.1_ASM258097v1 | *B. toyonensis* |
| GCF_002581125.1_ASM258112v1 | *B. toyonensis* |
| GCF_002581205.1_ASM258120v1 | *B. toyonensis* |
| GCF_002581275.1_ASM258127v1 | *B. toyonensis* |
| GCF_002581325.1_ASM258132v1 | *B. toyonensis* |
| GCF_002581455.1_ASM258145v1 | *B. toyonensis* |
| GCF_002581485.1_ASM258148v1 | *B. toyonensis* |
| GCF_002581675.1_ASM258167v1 | *B. toyonensis* |
| GCF_002581735.1_ASM258173v1 | *B. toyonensis* |
| GCF_002581765.1_ASM258176v1 | *B. toyonensis* |
| GCF_002581925.1_ASM258192v1 | *B. toyonensis* |
| GCF_002581955.1_ASM258195v1 | *B. toyonensis* |
| GCF_002582025.1_ASM258202v1 | *B. toyonensis* |
| GCF_002582095.1_ASM258209v1 | *B. toyonensis* |
| GCF_002582205.1_ASM258220v1 | *B. toyonensis* |
| GCF_002582715.1_ASM258271v1 | *B. toyonensis* |
| GCF_002583715.1_ASM258371v1 | *B. toyonensis* |
| GCF_002585135.1_ASM258513v1 | *B. toyonensis* |
| GCF_002585395.1_ASM258539v1 | *B. toyonensis* |
| GCF_002585525.1_ASM258552v1 | *B. toyonensis* |
| GCF_002588635.1_ASM258863v1 | *B. toyonensis* |
| GCF_002588695.1_ASM258869v1 | *B. toyonensis* |
| GCF_002588835.1_ASM258883v1 | *B. toyonensis* |
| GCF_002588905.1_ASM258890v1 | *B. toyonensis* |
| GCF_002589195.1_ASM258919v1 | *B. toyonensis* |
| GCF_002589255.1_ASM258925v1 | *B. toyonensis* |
| GCF_002589295.1_ASM258929v1 | *B. toyonensis* |
| GCF_002589305.1_ASM258930v1 | *B. toyonensis* |
| GCF_002589375.1_ASM258937v1 | *B. toyonensis* |
| GCF_002589595.1_ASM258959v1 | *B. toyonensis* |
| GCF_002589605.1_ASM258960v1 | *B. toyonensis* |
| GCF_002813875.1_ASM281387v1 | *B. cereus* |
| GCF_002893805.1_ASM289380v1 | *B. subtilis* |
| GCF_002982175.1_ASM298217v1 | *B. subtilis* |
| GCF_003013315.1_ASM301331v1 | *B. cereus* |
| GCF_003020845.1_ASM302084v1 | *B. cereus* |
| GCF_003148355.1_ASM314835v2 | *B. subtilis* |
| GCF_003148415.1_ASM314841v1 | *B. subtilis* |
| GCF_003184225.1_ASM318422v1 | *B. subtilis* |
| GCF_003227955.1_ASM322795v1 | *B. anthracis* |
| GCF_003386775.1_ASM338677v1 | *B. mycoides* |
| GCF_003410255.1_ASM341025v1 | *B. anthracis* |
| GCF_003410355.1_ASM341035v1 | *B. anthracis* |
| GCF_003426125.1_ASM342612v1 | *B. subtilis* |
| GCF_003445395.1_ASM344539v1 | *B. thuringesis* |
| GCF_003546665.1_ASM354666v1 | *B. thuringesis* |
| GCF_003568565.1_ASM356856v1 | *B. cereus* |
| GCF_003610835.1_ASM361083v1 | *B. toyonensis* |
| GCF_003610955.1_ASM361095v1 | *B. subtilis* |
| GCF_003612735.1_ASM361273v1 | *B. subtilis* |
| GCF_003612955.1_ASM361295v1 | *B. mobilis* |
| GCF_003626955.1_ASM362695v1 | *B. thuringesis* |
| GCF_003665195.1_ASM366519v1 | *B. subtilis* |
| GCF_003665215.1_ASM366521v1 | *B. subtilis* |
| GCF_003665235.1_ASM366523v1 | *B. subtilis* |
| GCF_003665255.1_ASM366525v1 | *B. subtilis* |
| GCF_003665275.1_ASM366527v1 | *B. subtilis* |
| GCF_003665295.1_ASM366529v1 | *B. subtilis* |
| GCF_003665315.1_ASM366531v1 | *B. subtilis* |
| GCF_003665335.1_ASM366533v1 | *B. subtilis* |
| GCF_003665355.1_ASM366535v1 | *B. subtilis* |
| GCF_003665395.1_ASM366539v1 | *B. subtilis* |
| GCF_003858675.1_ASM385867v1 | *B. pacificus* |
| GCF_003966295.1_ASM396629v1 | *B. albus* |
| GCF_003991175.1_ASM399117v1 | *B. thuringesis* |
| GCF_004006495.1_ASM400649v1 | *B. cereus* |
| GCF_004023375.1_ASM402337v1 | *B. mycoides* |
| GCF_004101345.1_ASM410134v1 | *B. subtilis* |
| GCF_004101365.1_ASM410136v1 | *B. subtilis* |
| GCF_004101405.1_ASM410140v1 | *B. subtilis* |
| GCF_004101425.1_ASM410142v1 | *B. subtilis* |
| GCF_004101445.1_ASM410144v1 | *B. subtilis* |
| GCF_004101465.1_ASM410146v1 | *B. subtilis* |
| GCF_004101485.1_ASM410148v1 | *B. subtilis* |
| GCF_004101565.1_ASM410156v1 | *B. subtilis* |
| GCF_004101945.1_ASM410194v1 | *B. subtilis* |
| GCF_004103535.1_ASM410353v1 | *B. subtilis* |
| GCF_004103555.1_ASM410355v1 | *B. subtilis* |
| GCF_004103595.1_ASM410359v1 | *B. subtilis* |
| GCF_004119535.1_ASM411953v1 | *B. subtilis* |
| GCF_004119555.1_ASM411955v1 | *B. subtilis* |
| GCF_004119595.1_ASM411959v1 | *B. subtilis* |
| GCF_004119615.1_ASM411961v1 | *B. subtilis* |
| GCF_004119635.1_ASM411963v1 | *B. subtilis* |
| GCF_004119655.1_ASM411965v1 | *B. subtilis* |
| GCF_004119675.1_ASM411967v1 | *B. subtilis* |
| GCF_004119695.1_ASM411969v1 | *B. subtilis* |
| GCF_004119715.1_ASM411971v1 | *B. subtilis* |
| GCF_004119775.1_ASM411977v1 | *B. subtilis* |
| GCF_004119815.1_ASM411981v1 | *B. subtilis* |
| GCF_004119835.1_ASM411983v1 | *B. subtilis* |
| GCF_004119875.1_ASM411987v1 | *B. subtilis* |
| GCF_004328925.1_ASM432892v1 | *B. subtilis* |
| GCF_004771155.1_ASM477115v1 | *B. cereus* |
| GCF_004801195.1_ASM480119v1 | *B. cereus* |
| GCF_005153965.1_ASM515396v1 | *B. subtilis* |
| GCF_005155285.1_ASM515528v1 | *B. thuringesis* |
| GCF_005160425.1_ASM516042v1 | *B. subtilis* |
| GCF_005217685.1_ASM521768v1 | *B. mycoides* |
| GCF_005217805.1_ASM521780v1 | *B. mycoides* |
| GCF_005234095.1_ASM523409v1 | *B. subtilis* |
| GCF_005707595.1_ASM570759v1 | *B. cereus* |
| GCF_005849145.1_ASM584914v1 | *B. subtilis* |
| GCF_006088795.1_ASM608879v1 | *B. subtilis* |
| GCF_006088855.1_ASM608885v1 | *B. anthracis* |
| GCF_006094295.1_ASM609429v1 | *B. cereus* |
| GCF_006094475.1_ASM609447v1 | *B. subtilis* |
| GCF_006151925.1_ASM615192v1 | *B. thuringesis* |
| GCF_006165085.1_ASM616508v1 | *B. subtilis* |
| GCF_006349595.1_ASM634959v1 | *B. pacificus* |
| GCF_006349625.1_ASM634962v1 | *B. tropicus* |
| GCF_006349645.1_ASM634964v1 | *B. tropicus* |
| GCF_006364495.1_ASM636449v1 | *B. subtilis* |
| GCF_006384875.1_ASM638487v1 | *B. cereus* |
| GCF_006457285.1_ASM645728v1 | *B. tropicus* |
| GCF_006741845.1_ASM674184v1 | *B. subtilis* |
| GCF_006742565.1_ASM674256v1 | *B. anthracis* |
| GCF_007672275.1_ASM767227v1 | *B. tropicus* |
| GCF_007673305.1_ASM767330v1 | *B. mycoides* |
| GCF_007673655.1_ASM767365v1 | *B. mycoides* |
| GCF_007673715.1_ASM767371v1 | *B. mycoides* |
| GCF_007673755.1_ASM767375v1 | *B. toyonensis* |
| GCF_007674195.1_ASM767419v1 | *B. toyonensis* |
| GCF_007676425.1_ASM767642v1 | *B. tropicus* |
| GCF_007676595.1_ASM767659v1 | *B. nitratireducens* |
| GCF_007676605.1_ASM767660v1 | *B. nitratireducens* |
| GCF_007677835.1_ASM767783v1 | *B. mycoides* |
| GCF_007681065.1_ASM768106v1 | *B. nitratireducens* |
| GCF_007681185.1_ASM768118v1 | *B. mobilis* |
| GCF_007681195.1_ASM768119v1 | *B. mobilis* |
| GCF_007681365.1_ASM768136v1 | *B. nitratireducens* |
| GCF_007682005.1_ASM768200v1 | *B. paramycoides* |
| GCF_007682135.1_ASM768213v1 | *B. paranthracis* |
| GCF_007682155.1_ASM768215v1 | *B. paranthracis* |
| GCF_007682195.1_ASM768219v1 | *B. mobilis* |
| GCF_007682355.1_ASM768235v1 | *B. nitratireducens* |
| GCF_007682405.1_ASM768240v1 | *B. tropicus* |
| GCF_900094345.1_ASM90009434v1 | *B. mycoides* |
| GCF_900094655.1_ASM90009465v1 | *B. mycoides* |
| GCF_900094695.1_ASM90009469v1 | *B. mycoides* |
| GCF_900094915.1_ASM90009491v1 | *B. cytotox* |
| GCF_900094995.1_ASM90009499v1 | *B. mycoides* |
| GCF_900095005.1_ASM90009500v1 | *B. mycoides* |
| GCF_900095655.1_ASM90009565v1 | *B. cytotox* |

**Table S5** Sequence and origin of query proteins used to search for homo- and orthologs within subject genomes in Table S4 using parwise tBLASTn searches. EnaX_NM_Oslo: Query protein sequence is sequenced from amplified lab strain. EnaX_GCF_001044825: Query sequence protein is from the gene in the publicly available NVH 0095-75: GCF_001044825.1_ASM104482v1. NM 0095-75

| **PCR product** | |
| --- | --- |
| Ena1B_NM_Oslo | MGNCSTNLSCCANGQIIVQDKVCIDWTAAATAAIIYADNISQDIYASGYLKVDTGTGPVTIVFYSGGVTGTAVETIVVATGSSASFTVRRFDTVTILGTAAAETGEFCMTIRYTLS |
| **From the GCF_001044825.1:** | |
| Ena1A_GCF_001044825, KMP91697.1 | MACECSSTVLTCCSDNSSNFVQDKVCNPWSSAEASTFTVYANNVNQNIVGTGYLTYDVGPGVSPANQITVTVLDSGGGTIQTFLVNEGTSISFTFRRFNIIQITTPATPIGTYQGEFCITTRYLMA |
| Ena1C_GCF_001044825, KMP91699.1 | LKPHKNIGCFAPLSIICQPTCPCPPPILPPERGDAELVTNEFAGDILISNDFIPISQKQLKQTNTTVNIWKNDGIVSLSGTISIYNNRNSTNALSIQIISSTTNTFTALPGNTISYTGFDLQSVSVIDIPSDPSIYIEGRYCFQLTYCKSKRDCL |

**References**

Argimon, S., Abudahab, K., Goater, R.J.E., Fedosejev, A., Bhai, J., Glasner, C., Feil, E.J., Holden, M.T.G., Yeats, C.A., Grundmann, H.*, et al.* (2016). Microreact: visualizing and sharing data for genomic epidemiology and phylogeography. Microb Genom *2*, e000093.

Burnley, T., Palmer, C.M., and Winn, M. (2017). Recent developments in the CCP-EM software suite. Acta Crystallogr D Struct Biol *73*, 469-477.

Farabella, I., Vasishtan, D., Joseph, A.P., Pandurangan, A.P., Sahota, H., and Topf, M. (2015). TEMPy: a Python library for assessment of three-dimensional electron microscopy density fits. J Appl Crystallogr *48*, 1314-1323.

Price, M.N., Dehal, P.S., and Arkin, A.P. (2010). FastTree 2--approximately maximum-likelihood trees for large alignments. PLoS One *5*, e9490.

Terwilliger, T.C., Ludtke, S.J., Read, R.J., Adams, P.D., and Afonine, P.V. (2019). Improvement of cryo-EM maps by density modification. bioRxiv.
